# Structure and Activation Mechanism of a Lamassu Phage Defence System

**DOI:** 10.1101/2025.03.14.643221

**Authors:** Yan Li, David W. Adams, Hon Wing Liu, Steven J. Shaw, Emiko Uchikawa, Milena Jaskólska, Sandrine Stutzmann, Laurie Righi, Mark D. Szczelkun, Melanie Blokesch, Stephan Gruber

## Abstract

Lamassu is a diverse family of defence systems that protect bacteria, including pandemic strains of *Vibrio cholerae*, against phage infection. They target essential cellular processes, aborting infection and preventing phage propagation by terminating the infected host. The mechanisms by which Lamassu effectors are activated when needed and otherwise suppressed are unknown. Here, we present structures of a Lamassu defence system from *Salmonella enterica*. We show that an oligomerization domain of the nuclease effector, LmuA, is sequestered by two tightly-folded SMC-like LmuB protomers and LmuC. Upon activation, liberated LmuA proteins assemble into a cyclic homo-tetramer, in which two of four nuclease domains are brought into proximity to create an active site capable of cleaving DNA. We propose tetramer formation is likely a one-way switch that establishes a threshold to limit potential spontaneous activation and cell death. Our findings reveal a mechanism of cellular defence, involving liberation and oligomerization of immune effectors, and shed light on how Lamassu systems balance potent immune responses with self-preservation.

## Introduction

Mobile genetic elements (MGEs) such as phages and plasmids exploit cellular resources to proliferate. To counteract MGEs, bacteria have evolved a wide array of defences ^1–3^. On bacterial genomes, defence systems tend to cluster together into genomic islands (‘defence islands’) and closely related genomes often harbour complementary sets of defence systems. These characteristics have led to the rapid identification of many new bacterial defence systems ^4–8^. Among these is a diverse family collectively termed Lamassu (after an Assyrian protective deity) which can be found in about 10% of bacterial and archaeal genomes ^7^. Initially, Lamassu systems were described as two-part modules, comprising a predicted nuclease effector, LmuA, and a putative DNA sensor, LmuB. Subsequent discoveries revealed a large variety of enzymatically diverse LmuA effectors and the presence of an additional ‘middle component’, LmuC, in most Lamassu systems ^7,9^, being encoded between LmuA and LmuB completing the Lamassu LmuACB operon (also denoted as DNA-defence modules DdmABC ^10^ or ATP-binding cassette-three component systems ‘ABC-3C’ ^11^).

The Lamassu protein LmuB is closely related to the SMC-like protein Rad50 ^11^. Canonical SMC proteins form multi-subunit ATP-binding cassette (ABC)-ATPase DNA motors that promote DNA loop extrusion for genome folding (cohesin, condensin, prokaryotic SMC) and extra-chromosomal circular DNA restriction (Smc5/6, Wadjet) ^12,13^. Rad50, together with its nuclease partner Mre11, prepares DNA double strand breaks for repair via homologous recombination and non-homologous end-joining ^14–16^.

Rad50 and LmuB proteins harbour an ABC head domain, a coiled coil emanating from the head, and a Zinc-hook (‘Zh’) dimerization domain at the other end of the coiled coil (Fig. 1C; Extended Data Fig. 1A). Like SMC proteins, Rad50 dimers undergo a rod-shaped to ring-shaped conformational switch upon ATP-mediated engagement of the two head domains (‘head-engagement’) ^17,18^ (Extended Data Fig. 1A). ATP-engaged heads expose a DNA binding surface between the two coiled coils, allowing the ring-shaped Rad50 dimer to engage various types of DNA ends including free and protein-blocked ends ^19,20^. LmuB proteins harbour a variant Walker B motif in which a catalytically important glutamate is replaced by glutamine; a substitution known to hinder ATP hydrolysis when introduced into other ABC ATPases including Rad50 and SMC proteins ^10,11,21^ (Fig. 1C).

**Fig. 1.**
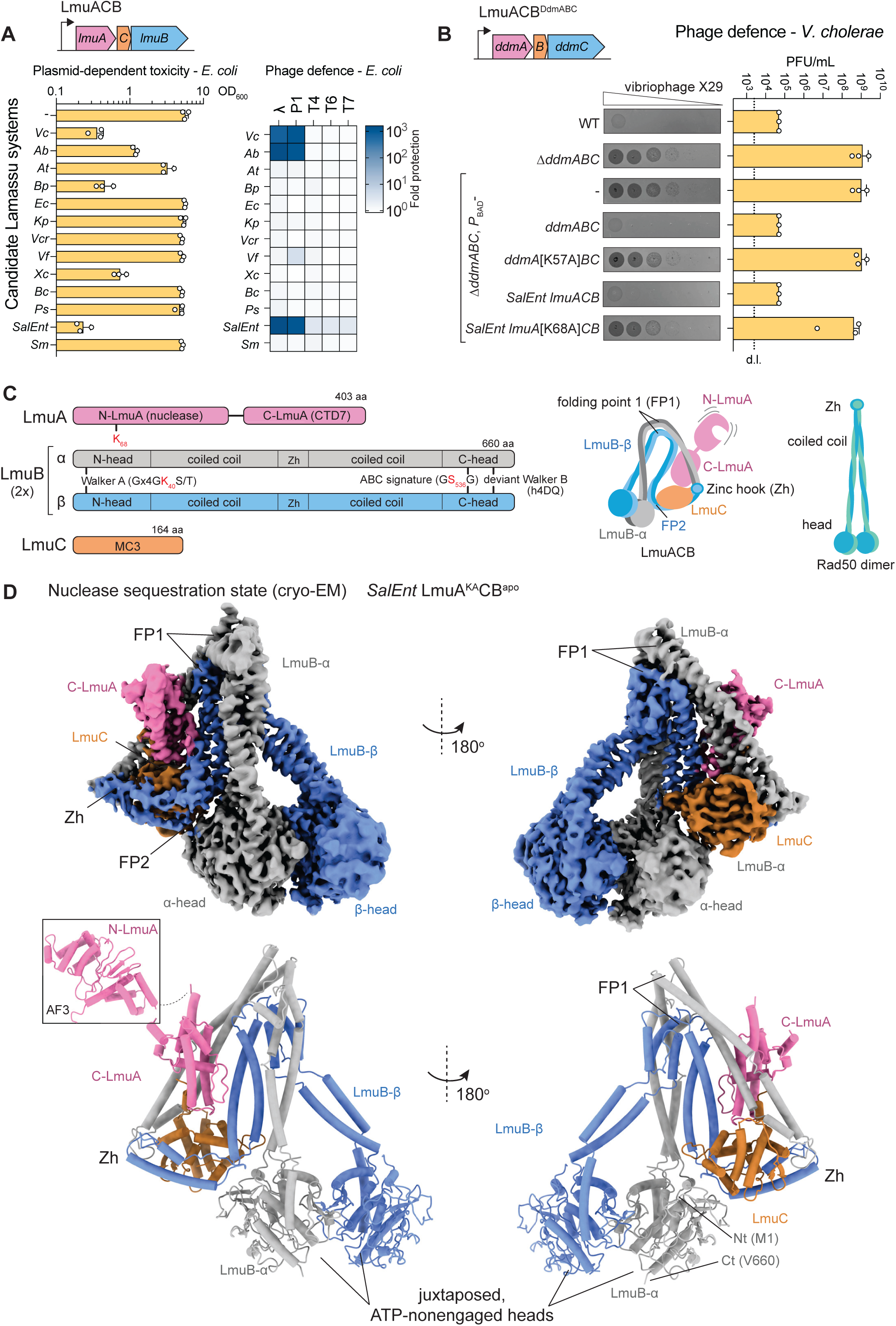
Structure of LmuACB^apo^ reveals a nuclease sequestration state. (A) Preselection of Lamassu systems based on plasmid-dependent toxicity (left panel) and phage defence (right panel) in *E. coli*. Selected operons were cloned under the arabinose-inducible *P*BAD promoter and expressed from a plasmid (left) or inserted into the chromosome (right). The optical density (OD600) of cultures of plasmid-harbouring strains grown for 6 h in LB supplemented with arabinose was measured. Data points from three replicates, the mean, and the standard deviation from the mean are shown (left panel). Protection against a panel of coliphages was tested by agar overlay plaques assays (right panel) with *E. coli* strains in which the *lmuACB*-carrying transposons were integrated into the chromosome via Tn7 transposition. The colour code indicates the fold protection levels relative to the *E. coli* strain lacking the *lmuACB* construct. *Vc, Vibrio cholerae* (*lmuACB^ddmABC^* cluster in this organism); *Ab, Acinetobacter baumannii; At, Agrobacterium tumefaciens; Bp, Burkholderia pseudomallei; Ec, Escherichia coli; Kp, Klebsiella pneumoniae; Vcr, Vibrio crassostreae; Vf, Vibrio fischeri; Xc, Xanthomonas campestris; Bc, Burkholderia cepacia; Ps Pseudomonas syringae; SalEnt, Salmonella enterica; Sm, Sinorhizobium meliloti* (Table S1). (B) Phage defence in *V. cholerae*. Selected Lamassu operons were cloned under the *P*BAD promoter and integrated into the chromosome of a *lmuACB^ddmABC^*deletion strain by Tn7 transposon insertion. Protection against vibriophage X29 was assessed using agar overlay plaques assays (images) with plaque forming units (PFU) quantified on the right. Data points from three replicates, the mean, and the standard deviation from the mean are shown. The dotted line denotes the detection limit (d.l.). (C) Left panel: Domain structure of the four LmuACB subunits in the nuclease sequestration state: LmuA, LmuC, and two protomers of LmuB. Conserved motifs are denoted; red labels indicate catalytic residues. Right panels: Schematic representation of the LmuACB complex in the nuclease sequestration state and for comparison, the Rad50 dimer (right). (D) Cryo-EM structure of *SalEnt* LmuACB in the nuclease sequestration state. Electron density maps (top panels) and structural models in cartoon representation (bottom panels) in back view (left panels) and front view (right panels). Density and cartoons for the nuclease subunit LmuA, the LmuB-a subunit, the LmuB-1 subunit, and the LmuC subunit are shown in pink, grey, blue, and orange colours, respectively. The LmuB dimer with juxtaposed heads features multiple coiled coil folding points (FP1, FP2), engaging the carboxyterminal domain of LmuA as well as LmuC at the Zinc hook region.

While the putative LmuB DNA sensor has a conserved domain organization between Lamassu systems, LmuA and LmuC exhibit greater variability. LmuA harbours several domains; at the amino terminus it contains usually one, sometimes two putative effector domains with various predicted enzymatic activities (such as nuclease, protease, alpha/beta hydrolase, and dehydrogenase domains) ^7,11^. The carboxy-terminal domain also falls into various types (denoted as CTD1 to CTD12); some types associate with a specific effector, while others appear more promiscuous existing in conjunction with different types of effectors ^11^. LmuC is a small, single-domain protein that has been grouped into eight distinct types (from MC1 to MC8) based on the fold and patterns of conserved residues. How the LmuB DNA sensors cooperates with such a variety of LmuA and LmuC proteins is unclear.

The Lamassu system DdmABC in the seventh-pandemic strains of *V. cholerae* El Tor, here also denoted as LmuACB^DdmABC^ (comprising LmuA protein DdmA, LmuC protein DdmB, and LmuB protein DdmC), was first identified due to its ability to restrict plasmids ^10^. In its indigenous host bacterium, LmuACB^DdmABC^ facilitates plasmid restriction together with the prokaryotic Argonaute (pAgo) system DdmDE ^10,22^. LmuACB^DdmABC^ also exhibits activity against specific vibriophages, and against coliphages when expressed in *E. coli* ^10,23,24^. In general, Lamassu systems appear to protect bacterial populations by abortive infection (induced cell death) ^7,10^, implying that Lamassu effectors inactivate an essential component of the infected cell to prevent further spreading of the MGE. Conversely, this requires that Lamassu systems have their effectors repressed to permit normal cell proliferation.

The folding of individual Lamassu subunits has been probed by structure prediction ^10,24^. However, the overall architecture, including subunit composition and the protein-protein interfaces forming the holocomplex remain unclear. Additionally, it is unknown how the LmuA effectors are kept inactive in the absence of Lamassu targets and how LmuB may recognize these targets and activate the diverse LmuA effectors. Here, we present structural and biochemical findings that proposes that effector sequestration and oligomerization upon release are a central principle for the control of Lamassu effectors.

## Results

### Identification of Lamassu systems with anti-plasmid and anti-phage activity

To identify Lamassu systems suitable for structural and biochemical studies, we focused on candidates related to LmuACB^DdmABC^ from *V. cholerae* El Tor ^10^ with selected *lmuA* and *lmuC* genes also belonging to the NTD^Cap^^4^-CTD7 and MC3 family, respectively ^11^ (Table S1) (Fig. 1A). Several candidates exhibited toxicity when expressed from a plasmid in *E. coli* (Fig. 1A), an activity previously reported for *lmuACB^ddmABC^* ^10^. As for LmuACB^DdmABC^, some of these systems also provided protection against phage P1 and A (but not T4, T6, and T7) when expressed from the chromosome of *E. coli* (Fig. 1A). To avoid host toxicity during recombinant protein expression, we mutated the active site lysine residue in the PD-(D/E)xK nuclease domain of LmuA in the candidate systems to alanine (‘KA’) (Fig. 1C). We selected LmuACB derived from *Salmonella enterica subsp. enterica* serovar Tennessee (strain TXSC_TXSC08-19 Genbank accession CP007505; ^25^) (short ‘*SalEnt’*) for further characterization, due to robust and homogeneous protein complex expression of the catalytically dead LmuA(K68A), LmuA^KA^ variant (Extended Data Fig. 1B). The purified protein complexes did not hydrolyse ATP, but we detected low levels of ATP hydrolysis (∼1/min) in the presence of DNA, which is generally known to stimulate ATP hydrolysis by SMC complexes, despite LmuB’s deviant Walker B motif. This activity was abrogated in the LmuB ATP binding (K40I) and head engagement (S536R) mutant (Extended Data Fig. 1C), indicating LmuACB are active if inefficient ATP hydrolases. We were also able to purify wild-type *SalEnt* LmuACB, however, at reduced yields, presumably due to self-restriction (Extended Data Fig. 1B, left panel).

To establish whether *SalEnt* LmuACB performs physiological immune functions we tested its activity in various contexts. *SalEnt* LmuACB was active in phage defence when expressed in *E. coli* (Fig. 1A). It also conferred protection to a *V. cholerae* strain against infection by phage X29 (Fig. 1B), a vibriophage previously reported to be restricted by LmuACB^DdmABC^ ^26^. Given the comparable phage protection levels of both native and ectopically expressed LmuACB^DdmABC^ and *SalEnt* LmuACB (Fig. 1B), we used this anti-phage assay to evaluate the function of various *SalEnt* LmuACB mutants in *V. cholerae*. We thereby established that protection against phage X29 required a functional LmuA nuclease, LmuB zinc hook, and LmuB ATP binding and head-engagement motifs (Extended Data Fig. 1D). Equipped with a recombinant protein reconstitution system and functional assays, we then sought to establish the mechanism underlying *SalEnt* LmuACB immune defense in molecular detail.

### Structure of DNA-free Lamassu LmuACB

To determine the structure of Lamassu, we collected and processed cryo-electron microscopy (cryo-EM) data with purified *SalEnt* LmuA^KA^CB protein, initially in the absence of DNA ligands but in buffer supplied with 1 mM ATP. Single-particle reconstruction produced a density map at a nominal resolution of 3.21 Å, harbouring two protomers of LmuB and single copies of LmuA and LmuC (Fig. 1D, Extended Data Fig. 7A, 8Ai). All domains of LmuA^KA^CB were resolved except for the amino-terminal domain of LmuA (N-LmuA), which is flexibly connected to the carboxy-terminal domain (C-LmuA) (Fig. 1D) and was presumably too mobile to resolve.

Substantial folding of the LmuB coiled coils results in a compact structure. The two LmuB protomers, LmuB-a and LmuB-1, are polymorphic and exhibit distinct folds in their coiled coil regions, producing tightly interconnected LmuB proteins with an asymmetric overall architecture. The two LmuB head domains are juxtaposed, but ATP-nonengaged. We did not observe density corresponding to ATP in the predicted active sites, despite the presence of ATP during grid preparation (Extended Data Fig. 1E, 8Aii). The zinc hook motifs of the two LmuB protomers are closely juxtaposed. Due to lower local map resolution the presence of a zinc atom cannot be clearly discerned. In LmuB-a, the coiled coil displays a single prominent folding point (‘FP1’) roughly halfway from the head (aa 266-276 & 416-432) (Fig. 1D, Extended Data Fig. 8Aiii-iv). In LmuB-1, in addition to FP1, there is another folding point (‘FP2’), located about three-quarters of the way from the head (aa 321-335 & 379-383) (Fig. 1D, Extended Data Fig. 8Avii). The LmuA and LmuC subunits are in the vicinity of the zinc hook motifs, making extensive contacts with both LmuB subunits and with each other. The LmuC subunit adopts a central position, contacting C-LmuA to the zinc hook-proximal LmuB coiled coils and the LmuB-a head domain. Thus, LmuC appears to stabilize the compact configuration of LmuA^KA^CB by establishing a network of inter-domain protein-protein contacts (Extended Data Fig. 8Av-vii). To interrogate the relevance of protein-protein contacts within LmuACB, we mutated a distinct series of key interfacial residues and observed variable phenotypes ranging from loss-of-function to hyperactivity, which will be discussed below.

Since the LmuA^KA^CB structure was generated in the absence of a DNA substrate and lacks bound ATP, we hypothesize that it corresponds to a ‘nuclease sequestration’ state in which the nuclease is inactive. Although the nuclease domain is loosely tethered to the remainder of LmuACB and apparently exposed to contact DNA, given its structural similarity to other nucleases that require dimerization for DNA cleavage, we speculated that LmuA might similarly require oligomerisation. In the context of LmuACB, the LmuA may therefore be locked as an inactive monomer Homodimers are often observed in Type II restriction enzymes, and oligomerization has been proposed for the CBASS-associated nuclease Cap4, another PD-(D/E)xK family nuclease ^27^. In further support of this model, the putative oligomerisation domain of LmuA forms embedded contacts with LmuCB that we later determined (see below) are sterically incompatible with homomer formation and thereby interfere with activation.

### DNA substrate binding by Lamassu

Having determined the structure of LmuACB^apo^, we then sought to establish the mechanism of DNA binding. Given recent findings that hairpin-forming DNA sequences may activate Lamassu ^24^, we tested its binding to various fluorescein-labelled DNA substrates by fluorescence anisotropy ^28^. Purified *SalEnt* LmuA^KA^CB bound to a 40-mer DNA duplex with a dissociation constant of ∼180 nM, while binding to single-stranded DNA was comparatively weak (Fig. 2A, left). ATP was not required for DNA binding, suggesting LmuACB diverges from other SMCs in which ATP-mediated head engagement creates a major DNA binding surface. ATP addition instead decreased the apparent binding affinity for DNA, suggesting that ATP-head engagement may hinder or alter DNA association (Fig. 2A, right). Furthermore, we found that LmuA^KA^CB can protect DNA ends from degradation by T5 exonuclease (Fig. 2B); this protection is counteracted by the addition of ATP, substantiating the idea that DNA end binding is reduced upon ATP binding.

**Fig. 2.**
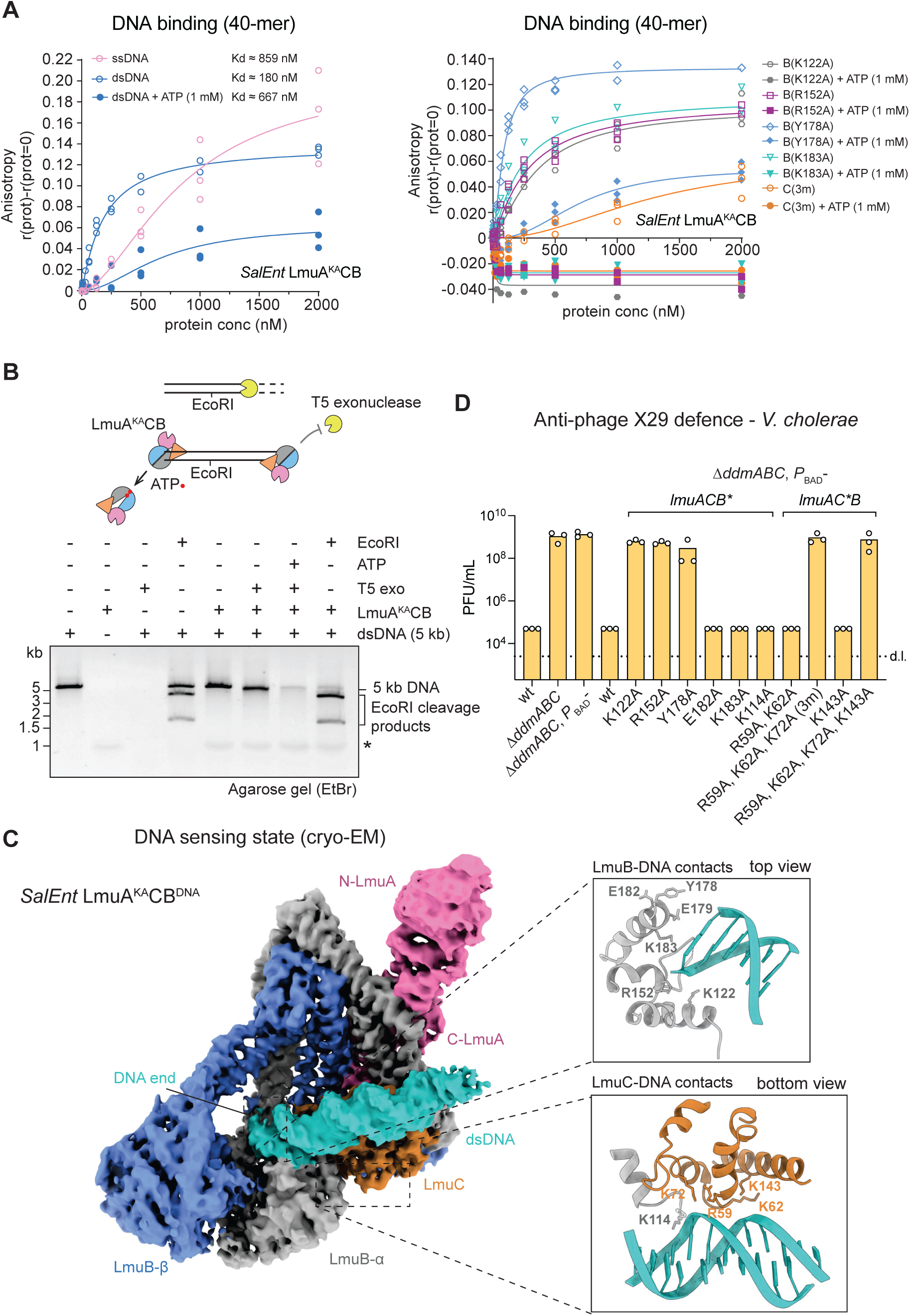
DNA end binding by LmuACB. (A) DNA binding measured by luorescence polarization with 6-FAM-labelled 40-mer single-stranded (ssDNA) and double-stranded (dsDNA) DNA with and without ATP (1 mM). The difference in anisotropy to the no-protein sample are plotted against the LmuA^KA^CB protein concentration (left graph). DNA binding for variants of the LmuA^KA^CB complex with and without ATP (1 mM) (right graph). Representative curves are shown. Estimates for the dissociation constants are given. Data points from three replicates. (B) DNA end protection from T5 exonuclease degradation by LmuA^KA^CB. Top panel: Illustration of the DNA end protection assay. Bottom panel: Agarose gel analysis of reactions with various enzymes as indicated on 5 kb linear DNA. LmuA^KA^CB blocks DNA degradation by T5 exo but not DNA cleavage by EcoRI. ATP addition alleviates DNA protection. A representative gel image is shown. The asterisk marks a band specific to the LmuA^KA^CB protein preparation. (C) Structure of the DNA sensing state of LmuA^KA^CB. Left panel: Cryo-EM map of the LmuA^KA^CB complex bound to a DNA end in front view. Right, top panel: LmuB contacts with the DNA end: residues selected for mutagenesis are highlighted. Bottom panel: LmuC contacts with internal DNA. (D) Selected DNA-binding mutants exhibit defects in phage defence. Phage defence against phage X29 in *V. cholerae* as measured by agar overlay plaque assay in triplicate. Details as described for Fig. 1B but using putative DNA-binding mutants of LmuB and LmuC as indicated. Data points from three replicates, the mean, and the standard deviation from the mean are shown. The dotted line denotes the detection limit (d.l.).

We next determined the structure of *SalEnt* LmuA^KA^CB bound to dsDNA by cryo-EM. We first purified stable complexes of LmuA^KA^CB with a 40-mer DNA duplex by size-exclusion chromatography, applied this sample to cryo-EM grids, and processed the resultant datasets (Extended Data Fig. 2A). We determined maps of LmuA^KA^CB bound to a DNA duplex (40 bp) at a nominal overall resolution of 3.08 Å (Fig. 2C, Extended Data Fig. 7B, 8Bi). The structure revealed LmuA^KA^CB bound to the end of the DNA ligand, in a manner distinct from previously characterized SMC-DNA complexes ^19,20^. Instead of DNA binding on top of ATP-engaged LmuB heads, the DNA approaches ATP-nonengaged heads from the side, with the DNA end primarily interacting with the LmuB-a head and with additional internal DNA contacts made with LmuC (Fig. 2C, Extended Data Fig. 8Bv-viii). This structure implies that Lamassu may recognize DNA ends by directly binding them, as opposed to sliding onto neighboring (end-proximal) DNA proposed for related Rad50/Mre11 complexes^18^. This distinction might be relevant for the recognition of the cognate DNA substrate by Lamassu. Notably, presence of ATP-nonengaged heads implies that head engagement is unfavorable or slow even in the presence of a DNA duplex as well as ATP (1 mM).

LmuA^KA^CB remained mostly unchanged in LmuA^KA^CB^DNA^ when compared to LmuA^KA^CB^apo^, with the notable exception of a somewhat increased visibility of N-LmuA and a slight shift in the coiled coil position (Fig. 2C, Extended Data Fig. 2B). We speculate that this conformation represents a ‘DNA sensing state’ rather than DNA cleavage state with the nuclease domain remaining distant from DNA. Accordingly, the DNA in this structure might correspond to a bound substrate that activates LmuACB, rather than the target DNA that is degraded by LmuACB.

Based on the LmuA^KA^CB^DNA^ structure, we mutated putative DNA binding residues on LmuB and LmuC (Fig. 2C). Individual point mutations on LmuB, K122A, R152A, or K183A, mildly reduced DNA binding by fluorescence anisotropy (Fig. 2A. right); K122A and R152A (but not K183A) also led to sensitivity to phage X29 infection *in vivo* (Fig. 2D). Since these LmuB residues may also contribute to DNA binding by ATP-engaged heads, we cannot exclude the possibility that the observed phenotypes result from loss of internal DNA binding rather than loss of DNA end binding. This possibility is supported by the observation that the K122A, R152A, and K183A mutations impact DNA binding also in the presence of ATP (Fig. 2A. right). Single mutations on LmuC failed to give appreciable phenotypes. However, an LmuC ‘3m’ variant (R59A, K62A, K72A) blocked DNA binding and also failed to provide phage protection in *V. cholerae* (Fig. 2B, 2D), indicating that DNA binding by LmuC is critical. Notably, another LmuB mutation, Y178A, had little impact on DNA binding in presence or absence of ATP (Fig. 2A. right), but lost *in vivo* phage protection (Fig. 2D). The side chain is located on top of the LmuB head domains and may promote or hinder head engagement or DNA binding to engaged heads.

### Lamassu DNA cleavage and its structural basis

Having established the structure of two different functional states of LmuACB, corresponding to putative nuclease sequestration and DNA sensing states, we next wanted to address the mechanism by which the LmuA nuclease becomes competent to cleave DNA substrates.

Given the structural similarity of LmuA with Type II restriction enzymes, in which a dimeric nuclease cleaves a single dsDNA substrate, we performed AlphaFold3 (AF3) predictions of two protomers of LmuA together with dsDNA. AF3 predicted a tightly associated dimer of N-LmuA nuclease domains bound to a DNA duplex (Fig. 3A, left panels); a similar architecture was observed with predictions for N-LmuA^DdmA^ (Extended Data Fig. 3A). In the predicted dimers, two active sites are positioned relative to each other in a configuration capable of generating DNA ends with a single-nucleotide 3’ overhang (Fig. 3A)—distinct from the four-nucleotide 5’ overhang produced by the restriction enzyme HindIII, another PD-(D/E)xK family member (PDB: 3WVG) (Fig. 3A, Extended Data Fig. 3B). The AF3 predictions suggest firstly that DNA cleavage may produce DNA ends with distinctive characteristics, specifically single-nucleotide 3’ overhangs. Secondly, LmuA activation may involve dimerization or oligomerization for nuclease activation, conceivably following its liberation from the larger LmuACB complex, in which the putative C-LmuA oligomerisation domain is otherwise occluded by contacts with LmuCB.

**Fig. 3.**
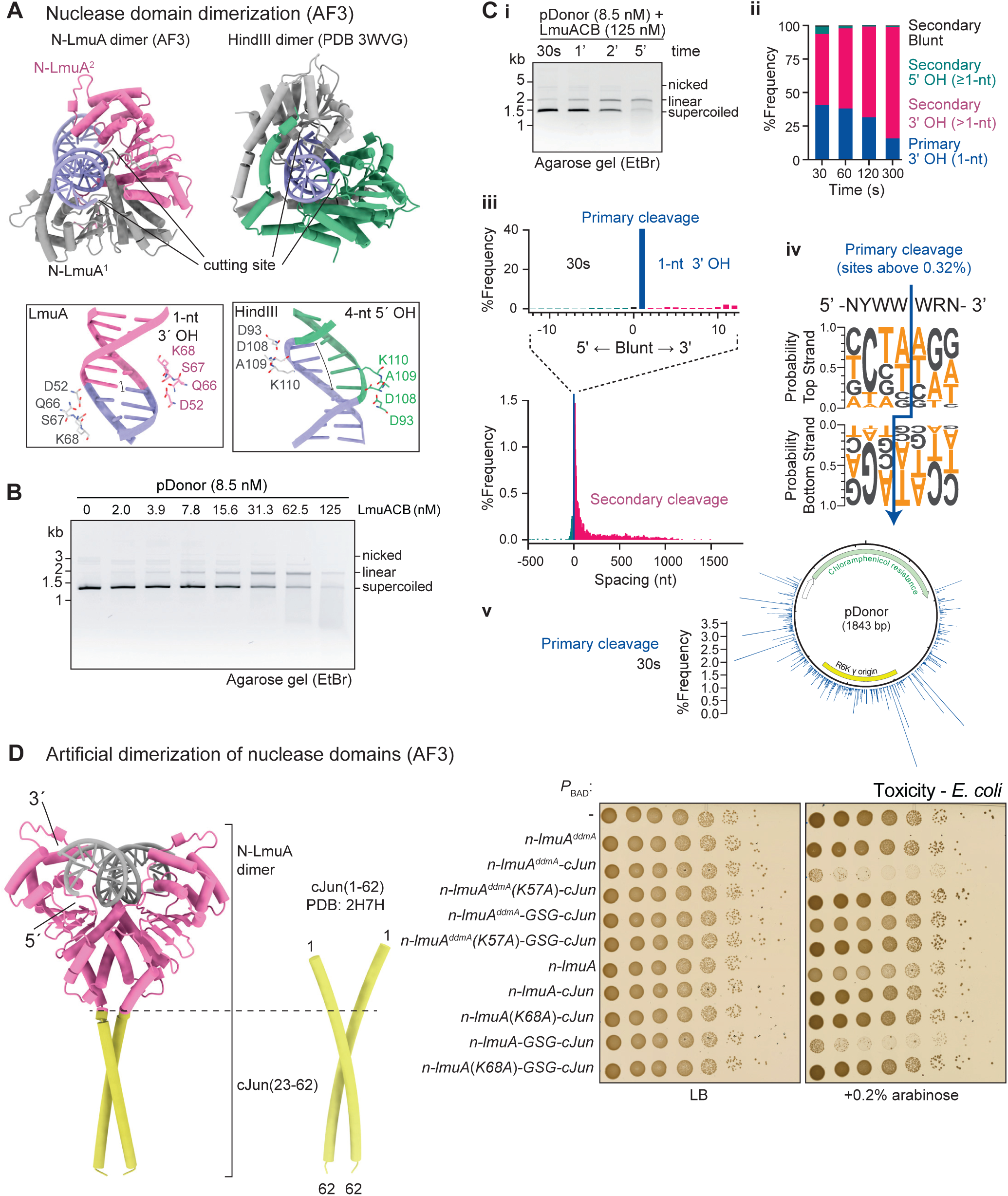
DNA cleavage by wild-type LmuACB. (A) AlphaFold3 (AF3) prediction of a DNA bound N-LmuA dimer (left) and its direct comparison to the structure of the restriction enzyme HindIII bound to DNA (PDB 3WVG, right). Note that one chain is shown in colour (pink and green for LmuA and HindIII, respectively) and the other chain in grey colours. The sites of DNA cleavage and the resulting overhangs for HindIII and LmuA as deduced from the positioning of the active site residues are indicated (corresponding bottom panels). (B) Lamassu DNA cleavage as analysed by agarose gel electrophoresis using supercoiled 1.8 kb pDonor as DNA substrate and increasing amounts of LmuACB in presence of ATP (1 mM). (C) Analysis of the LmuACB DNA cleavage pattern by ENDO-Pore sequencing. **i**, Time course of DNA cleavage to optimize yield of single cleavage events analyzed by agarose gel electrophoresis. **ii**, Proportion of linear DNA cleavage at each time point. **iii**, Frequencies of DNA end types (blunt, 3’ overhang or 5’ overhang) after 30 s incubation with LmuACB as determined by ENDO-Pore sequencing. Primary cleavage events (from a single cleavage event) are depicted in blue colours, secondary cleavage events depicted in red colours. Equivalent graphs for the later time points are shown in Extended Data Fig. 4. **iv**, DNA cleavage site preference obtained by ENDO-Pore sequencing for primary cleavage events with an occurrence of more than 0.32% (11.5% total, 19 sites). **v**, Cleavage site distribution on pDonor after 30 s incubation with LmuACB as determined by ENDO-Pore sequencing. Note that cleavage sites at the chloramphenicol resistance gene were not recorded as the gene was essential for generating the pUC-*ori* ligation library for ENDO-Pore (see methods). See Extended Data Fig. 4 and Extended Data Fig. 5 for further analyses of LmuACB-generated DNA end characterization with ENDO-Pore sequencing. (D) Generation of active N-LmuA dimers by artificial dimerization using cJun fusion proteins. Left panel: AF3 predictions of a selected N-LmuA-cJun fusion dimer with a DNA duplex. For reference, the structure of the corresponding cJun leucine zipper dimer is shown (PDB: 2H7H). Right panels: Toxicity of N-LmuA^DdmA^ and N-LmuA fusions to cJun when expressed in *E. coli* without (left) or with (right) induction by arabinose. The spots represent ten-fold serial dilutions from left to right.

To investigate the nature of DNA ends produced by Lamassu, we assessed the deoxy-ribonucleolytic activity of purified wild-type *SalEnt* LmuACB on a 1.8 kb test DNA substrate (‘pDonor’) ^29^. When supercoiled, nicked or linear forms of pDonor were incubated with increasing concentrations of LmuACB, we observed DNA degradation, resulting in a DNA smear by gel electrophoresis (Fig. 3B, Extended Data Fig. 3C). LmuACB cleavage products are readily re-ligated by T4 ligase, as expected for ends with single-nucleotide overhangs (Extended Data Fig. 3D). The nuclease activity was abolished by the KA mutation in LmuA, yet DNA degradation did not require ATP, nor a functional signature motif for ATP-head engagement in LmuB (Extended Data Fig. 3E, F). While the signature motif mutation (S536R) did not affect the nucleolytic activity *in vitro*, it eliminated the anti-phage activity of LmuACB when expressed in *V. cholerae* (Extended Data Fig. 1D). This suggests that the observed nuclease activity does not require ATP-dependent head engagement.

We proceeded to characterize the generated DNA ends using ENDO-Pore sequencing ^30^. Briefly, pDonor was exposed to LmuACB, the resulting DNA fragments (Fig. 3Ci) were end-blunted and ligated to a DNA insert containing a *pir*-independent replication origin. *E. coli* was transformed with the ligation products and approximately 20,000 transformants per timepoint were harvested and pooled. A plasmid DNA library was then prepared and analyzed using single-molecule nanopore sequencing. Reads with full-length pDonor sequences, which represent DNA from a single LmuACB cleavage event (‘primary cleavage events’), revealed a strong bias for ends with a single-nucleotide 3’ overhang (30 s data in Fig. 3Cii, iii, other timepoints and additional data in Extended Data Fig. 4) consistent with the AF3 structure prediction (Fig. 3A). There was a notable preference for cleavage at 5’-YWW↓WR-3’ sequences (where Y = C or T, W = A or T and R = A or G) (Fig. 3Civ). Over time smaller DNA fragments, presumably produced by additional LmuACB cleavage events (‘secondary cleavage events‘) emerged, exhibiting a similar cleavage site preference, with DNA lengths appearing to be randomly distributed, implying independent cleavage events at random positions rather than progressive DNA degradation (Fig. 3Cii, iii, v, Extended Data Fig. 4, 5). The degenerate cleavage site preference is consistent with the idea that Lamassu represents an abortive infection system that once activated degrades any DNA in the vicinity of the LmuA oligomer including chromosomal DNA.

To test whether N-LmuA homodimerization is involved in Lamassu nuclease activation, we engineered chimeric proteins in which a leucine zipper dimer formed by a truncated cJun protein, fused to N-LmuA/N-LmuA^DdmA^ sequences, brings together two nuclease domains in a configuration resembling the AF3 predictions (Fig. 3D, left panel, Extended Data Fig. 3G) ^31^. The N-LmuA- and N-LmuA^DdmA^-cJun chimeric proteins were toxic when expressed in *E. coli* (Fig. 3D, right panel). Their toxicity was abrogated by the respective catalytic lysine (KA) mutation in N-LmuA and N-LmuA^DdmA^. Curiously, it was sensitive to how cJun was connected to N-LmuA and N-LmuA^DdmA^ (with only one out of two constructs each being toxic), suggesting that an appropriate dimer geometry is crucial for nuclease activity. Together, these findings suggest that dimerization of N-LmuA is sufficient for nuclease activation. Since LmuACB harbours only a single protomer of LmuA, we conclude that nuclease activation requires either oligomerization of LmuACB or, more likely, of LmuA following its liberation from the LmuACB complex.

### LmuA oligomerization upon liberation from LmuACB

Given the correlation between multimerization of LmuA and nuclease activity, we then wanted to understand the mechanistic basis by which oligomerisation leads to LmuA activation. AF3 predictions with multiple copies of C-LmuA showed that the oligomerization of C-LmuA (Fig. 4A) appeared incompatible with simultaneous formation of LmuACB complexes, as many C-LmuA residues involved in packing to LmuCB also participate in the predicted C-LmuA oligomerization contacts (Fig. 4A, 4B) and due to steric clashes between LmuACB and additional promoters of C-LmuA (Extended Data Fig. 2C). This suggests that formation of LmuACB complexes and LmuA oligomers might be mutually exclusive, implying that Lamassu activation requires the unlocking of LmuA from LmuACB complexes.

**Fig. 4.**
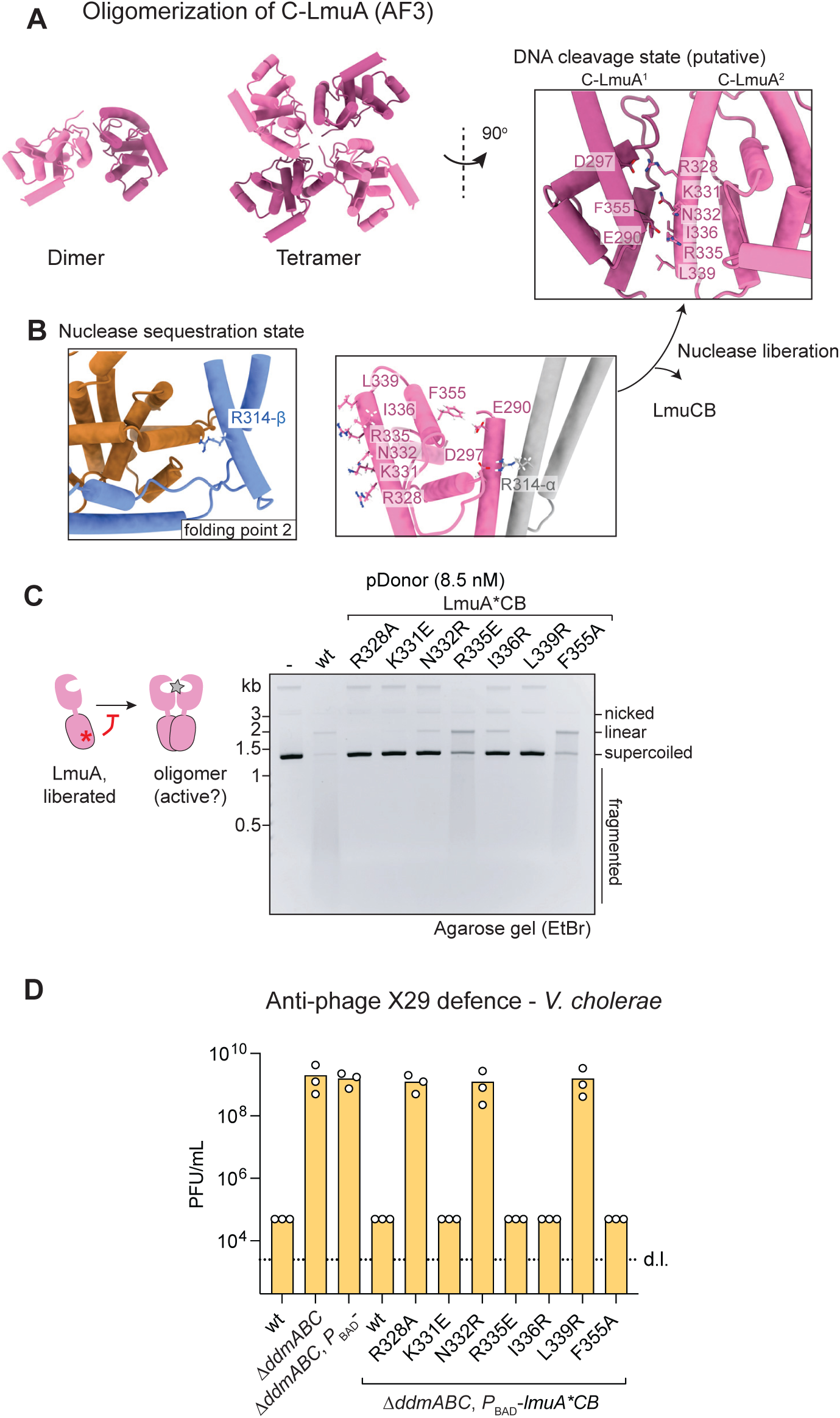
Oligomerization of the carboxyterminal domain of LmuA. (A) Selected AF3 predictions of multimeric forms of *SalEnt* C-LmuA shown in cartoon representation. (B) Selection of residues for mutagenesis to disrupt the C-LmuA oligomerization interface based on the AF3 tetramer model (A, right panel). The position of the equivalent residues in the nuclease sequestration state are denoted (left panels). Residues that contribute to the interactions in both states were excluded from the mutagenesis. (C) DNA cleavage activity of mutants of LmuACB putatively defective in C-LmuA oligomerization. Red asterisk represents C-LmuA mutations preventing C-LmuA oligomerization. Right panel: Agarose gel analysis of pDonor cleavage showing the impact of the indicated LmuA mutations on nucleic acid degradation. (D) Phage X29 defence in *V. cholerae* by LmuACB operons harbouring LmuA mutations. Details as described in Fig. 1B with LmuA mutations described in panel C. Data points from three replicates, the mean, and the standard deviation from the mean are shown. The dotted line denotes the detection limit (d.l.).

If oligomerization underpins LmuA activation, two key predictions arise: mutating the predicted C-LmuA oligomerization interface should result in nuclease-dead, loss-of-function phenotypes, and disrupting the C-LmuA interface with LmuC and LmuB should produce hyperactive mutants, due to a higher propensity for LmuA liberation. To test the first prediction, we targeted residues in C-LmuA that contribute to the predicted C-LmuA oligomerization interface but are not involved in contacting LmuB and LmuC (Fig. 4A, 4B). Individually mutating R328 to alanine, K331 to glutamate, and N332 and L339 to arginine indeed abolished nuclease activity, while substituting R335 with glutamate and F335 with alanine significantly reduced nuclease function (Fig. 4C). The R328A, N332R, and L339R mutations also inhibited phage defence by LmuACB in *V. cholerae* (Fig. 4D), suggesting that LmuA oligomerization is essential for phage defence by Lamassu. Two of the mutants (K331E and I336R) that were inactive *in vitro* remained functional for phage defence *in vivo*, indicating that oligomerization might be more robust *in vivo*, possibly owing to macromolecular crowding.

To test the second prediction, we targeted residues at the interface of C-LmuA with LmuB and LmuC but excluded residues from the analysis which apparently also contribute to C-LmuA oligomerization (Fig. 5A). Introducing LmuB^GR^ or LmuA^TR^, *i.e.* arginine substitutions of LmuB G318 and LmuA T306 respectively, shifted the elution of LmuACB proteins to a later volume in size exclusion chromatography, suggesting alterations in the architecture or composition of the mutant LmuACB complexes such as disintegration of the protein complex to produce smaller sub-complexes (Fig. 5B). In contrast, another mutation in the same interface, F303A in LmuA (‘FA’), did not show this shift in elution volume (Extended Data Fig. 6A). The LmuB^GR^ and LmuA^TR^ mutations, but not LmuA^FA^, resulted in dramatically increased nuclease activity *in vitro*, with a roughly more than a tenfold enhancement in DNA degradation rate (Fig. 5C), and in hyper-toxicity phenotypes *in vivo* when expressed in *V. cholerae* (Fig. 5D, Extended Data Fig. 6B). Mutating equivalent residues in LmuACB^DdmABC^ was also lethal in *V. cholerae* (Extended Data Fig. 6B). Together, these findings strongly support the idea that LmuA binding to LmuCB hinders activation, while its binding to other LmuA proteins promotes nuclease activity, consistent with the notion of nuclease inactivation by sequestration in LmuACB and subsequent liberation to form active LmuA oligomers.

**Fig. 5.**
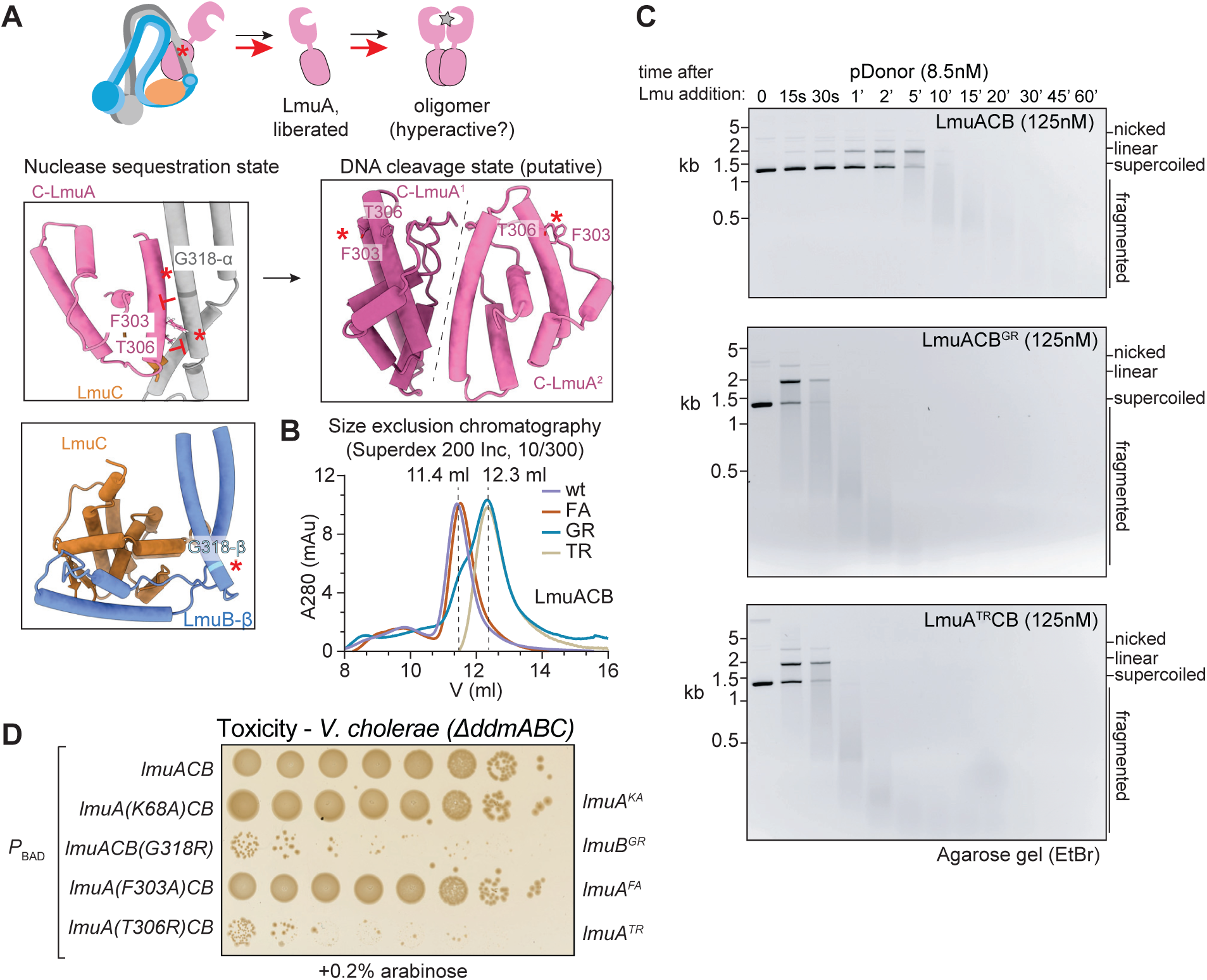
Mutational destabilization of LmuACB leads to hyperactivity and -toxicity. (A) Upper panel: Depiction of the mutagenesis strategy to promote LmuA liberation from sequestration by LmuACB. Lower panels: residues selected for mutagenesis to promote LmuA liberation from LmuACB in the nuclease sequestration state (left panels). The selected mutations are expected to not interfere with C-LmuA oligomerization (right panel). (B) Elution of LmuACB and variants thereof (F303A, FA; G318R, GR; T306R, TR) from a size exclusion chromatography column. The elution maxima shift from about 11.4 to 12.3 ml, indicating disintegration of the LmuACB complex. (C) DNA cleavage activity of mutant LmuACB on pDonor. DNA samples were taken at the indicated time points and analyzed by agarose gel electrophoresis. (D) Toxicity of mutant variants of LmuABC when expressed in *V. cholerae* with induction by arabinose, as described in Fig. 3D.

### Cryo-EM structure of the active Lamassu nuclease tetramer

Preliminary cryo-EM analysis of LmuA^KA^CB^GR^ indicated the presence of LmuA oligomers, along with other particles, potentially corresponding to LmuACB and LmuCB sub-complexes. Aiming to improve single particle reconstruction, we isolated pure preparations of LmuA^KA^ protein by separating it from LmuCB^GR^ and LmuA^KA^CB^GR^. This was achieved by including a high salt wash (with 1.5 M NaCl) during protein purification when LmuA^KA^CB^GR^ is immobilized on glutathione beads by the GST tag on LmuA. After further purifying and concentrating the LmuA^KA^ sample (Fig. 6A), we showed that it efficiently bound a 45-mer DNA duplex in electrophoretic mobility shift assays. We obtained well-distributed particles on grids using a chameleon® device ^32^, allowing sufficient particle picking for 3D reconstruction at an estimated resolution of 3.26 Å (Extended Data Fig. 7C). Cryo-EM analysis showed DNA-bound LmuA^KA^ tetramers (Fig. 6B, Extended Data Fig. 9). In the LmuA tetramer, two of the four N-LmuA interact with a single DNA duplex, while the third lacks DNA contact and the fourth is poorly resolved. The C-LmuA tetramer showed modest flexibility relative to the active site N-LmuA dimer, as indicated by the lower local resolution as well as by 3D-variability analysis (Extended Data Fig. 7C, movie 1). The active site residues are positioned to generate a single nucleotide 3’ overhang (Fig. 6C) as confirmed by ENDO-Pore sequencing. These findings are consistent with the idea that activation of Lamassu requires the release of LmuA proteins, presumably from multiple LmuACB complexes, to allow for the formation of an LmuA homo-oligomer that represents the DNA cleavage state of the Lamassu nuclease belonging to the LmuA CTD7 family.

**Fig. 6.**
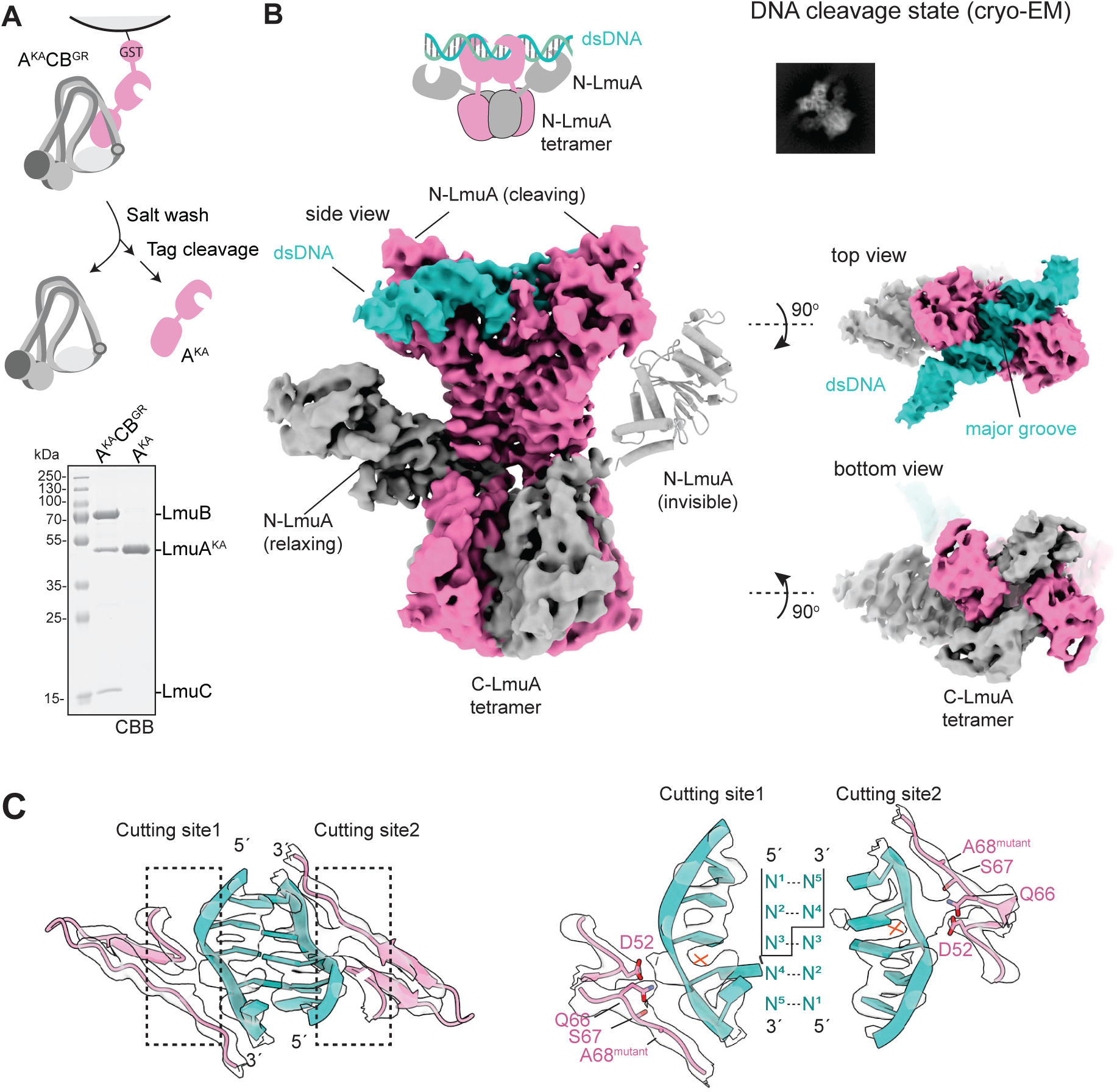
Liberated LmuA forms a homo-tetrameric DNA cleaving complex. (A) Cartoon illustrating the strategy for isolating LmuA^KA^ from LmuA^KA^CB^GR^ complexes during protein purification (top panels). SDS PAGE analysis of purified preparations of LmuA^KA^CB^GR^ and LmuA^KA^, confirming the presence or absence of LmuB and LmuC subunits. CBB, Coomassie Brilliant Blue staining. (B) Structure of the DNA-bound LmuA tetramer. Top left panel: Representative 2D class showing both cutting and relaxing N-LmuA domains. Top right panel: local cryo-EM density map indicating residues at the C-LmuA oligomerization interface. Bottom panels: Cryo-EM map of the tetrameric LmuA bound to a 45-mer DNA duplex in sideview; one N-LmuA domain is lacking in the electron density map; its putative position is indicated by the AF3 model. Top and bottom views are shown on the right (top and bottom panel, respectively). (C) The local electron density showing details of the DNA contact of the LmuA active site (cutting site 1&2) and a 1nt 3’-cutting end is indicated too (right panel).

## Discussion

Bacterial defence systems are widespread and diverse but for many, we do not understand how they are kept in check in a state that does not affect the host yet are efficiently unleashed by MGE presence. In this study, we shed light on molecular mechanism of one such system revealing a paradigm of activation through unlocking of the effector oligomerization domain (Fig. 7A). This paradigm may apply not only to members of the Lamassu^Cap^^4-CTD7-MC3^ sub-family, including *V. cholerae* LmuACB^DdmABC^ and *SalEnt* LmuACB, but could also extend to other sub-families with non-nuclease effectors, provided their activation depends on oligomerization (see below).

**Fig. 7.**
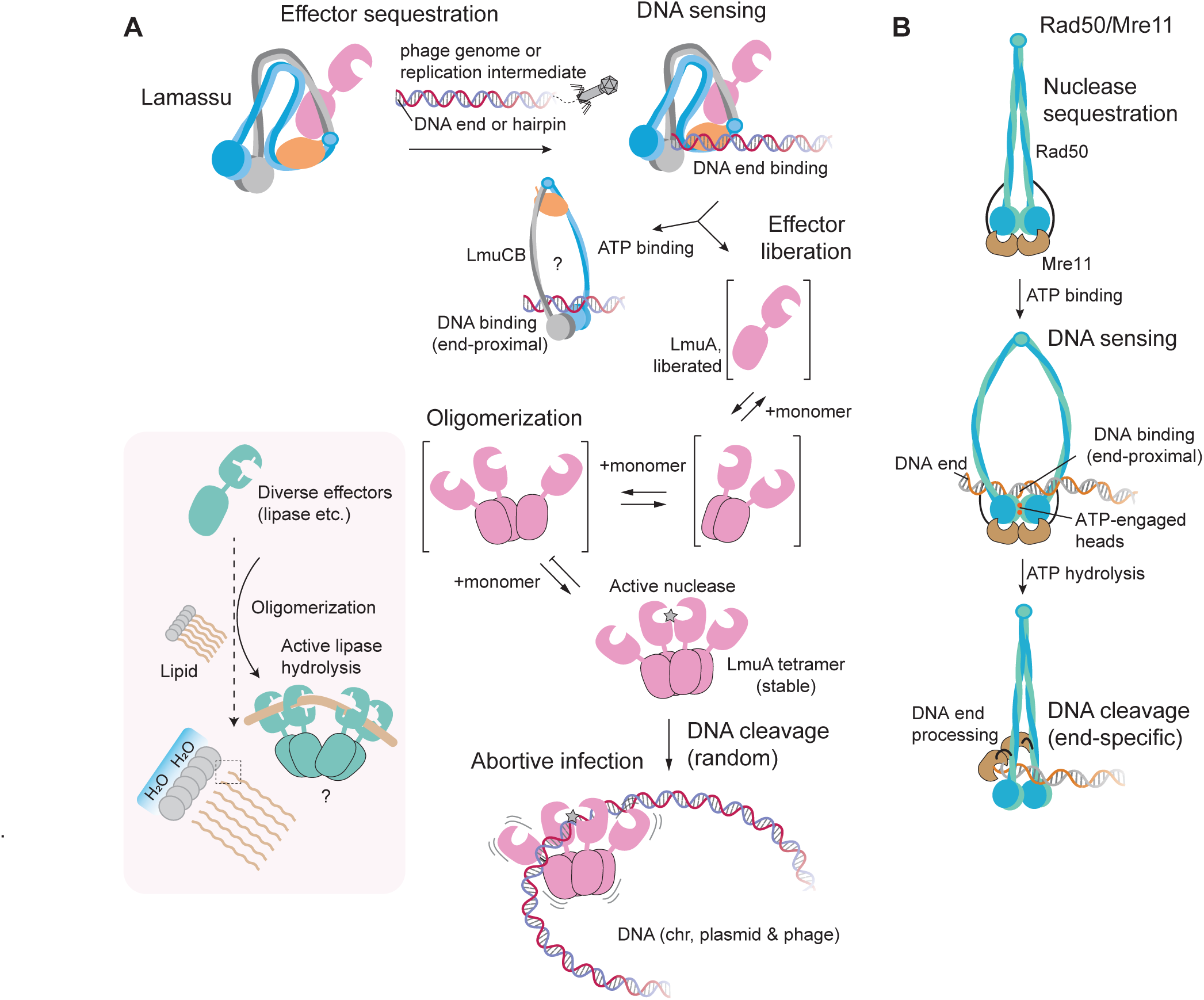
Model for Lamassu activation. (A) LmuACB complexes are formed from newly synthesized proteins. LmuA in isolation might not fold correctly and be nonfunctional (not shown). LmuACB complexes scan the cell for aberrant DNA secondary structures such as DNA ends and DNA hairpins. Upon DNA recognition involving DNA end binding, ATP-mediated head engagement converts LmuACB from the sequestration state to a putative open state, leading to the concomitant liberation of LmuA. Liberated LmuA undergoes oligomerization eventually producing LmuA tetramers, representing the active Lamassu effector nuclease for nucleic acid cleavage and abortive infection. Insert: Activation by liberation and oligomerization might apply to distant Lamassu relatives using other effectors such as lipases. (B) For comparison, a model of Rad50/Mre11 and its activation for DNA end processing is shown^18^.

Nucleases serve as effectors in diverse bacterial defence systems with the control of nucleic acid cleavage being achieved by two distinct principles: either the nuclease is permanently ‘on’ and the host DNA is enzymatically modified to prevent self-restriction (e.g. Restriction-Modification systems, BREX, DISARM) or the nuclease is produced in a neutralized ‘of’ state and only activated upon the detection of MGEs (e.g. Gabija, Paris, Hachiman, DdmDE, Wadjet) ^22,33–38^. Since accidental activation is fraught with severe, potentially lethal, consequences, activation often involves a threshold setting regulatory mechanism, limiting it to situations with sufficiently strong invader signal ^39^. We put forward a new mechanism for threshold signalling that works independent of second messenger modules and is based on the oligomerization of a rare form: the liberated effector protein.

### Neutralization by sequestration of effector oligomerization domain

We envisage that the compact architecture of Lamassu plays a key role in strictly controlling the effector, preventing its accidental activation in the absence of MGEs. This might be critical if activation is indeed nearly irreversible due to the inefficiency of LmuB in hydrolysing ATP and the putative relative stability of LmuA tetramers. We show that LmuACB forms a compact structure with an extensive interaction network involving the four subunits. C-LmuA contacts all three partner proteins, while N-LmuA fails to form visible contacts. The exclusivity of the protein-protein interactions to the C-LmuA oligomerization domain explains how an effector domain could be freely substituted for a structurally unrelated one without disrupting the regulatory mechanisms, providing flexibility and robustness allowing evolutionary change likely contributing to past and ongoing successes in the diversification of the Lamassu family of bacterial defence systems.

A curious observation in our study is that N-LmuA flips to the opposite side of C-LmuA between the sequestration state and the active state (Extended Data Fig. 6D, E). This conformational shift might serve as a safety feature, preventing a single liberated LmuA protein from prematurely forming an active dimer together with the LmuA protomer of an LmuACB complex, thereby ensuring that oligomerization—driven by sufficient liberated LmuA concentrations—is required for activation (Fig. 7A). This mechanism not only enhances the precision of Lamassu activation but also underscores its evolutionary refinement as a defence system. Whether this conformational change is a prerequisite for oligomerization and could thus explain why ‘free’ LmuA^DdmA^, produced from the chromosome in the absence of LmuC^DdmB^ and LmuB^DdmC^, exhibits relatively limited toxicity remains to be determined (albeit *SalEnt* LmuA exhibits toxicity when expressed on its own from a plasmid in *E. coli*) ^10^. For the unlocking and oligomerization model to function, newly synthesized ‘free’ LmuA must be nonfunctional, either by being mis-folded or unstable/degraded.

### Liberation and Oligomerization: A One-Way Switch

Many enzymes including nucleases are active in a dimeric form (e.g. restriction endonucleases, ABC transporters). Higher-order oligomers are also common, with all subunits typically contributing to the activity, as seen in AAA+ ATPase hexamers. In case of Lamassu, oligomerization might be important for strict regulation rather than activity itself. Since a single active LmuA nuclease is conceivably able to generate sufficient damage to the chromosome to kill the bacterial cell, its assembly must be avoided in a healthy cell. Assembly is likely inefficient due to the presence of minute concentrations of ‘liberated’ LmuA and the relatively high dissociation constant for an isolated C-LmuA oligomerization contact. In a cyclic oligomer (such as the tetramer), each subunit is held in place by oligomerization contacts with two neighbouring C-LmuA protomers, providing extra stability relative to smaller linear assemblies and representing a one-way switch (from of to on). The dynamics of the switch might be fine-tuned by the number of subunits forming the active oligomer (‘n-merisation’) with more dangerous effectors forming even larger assemblies. We indeed found LmuA proteins with predicted lipase effector domains, belonging to CTD8 and CTD9 family of LmuA proteins, capable of forming dimers and higher-order cyclic oligomers (but not cyclic tetramers) by AF3 prediction (Extended Data Fig. 6C). It remains to be established whether and how oligomerization of these non-nuclease effector domains leads to their activation.

LmuA oligomerization likely critically depends on the (local) protein concentration of liberated LmuA. This compares to the ligand-dependent oligomerization proposed for the CBASS-associated Cap4 nuclease, where the nucleotide ligand concentration presumably sets the threshold for oligomerization and activation ^27^. NLR immune receptors of the human innate immune system, including the NLRP3 or NLRC4 inflammasomes, also rely on ligand-induced oligomerization ^40^. Oligomerization is mediated by a central nucleotide-binding and oligomerization domain (the NACHT domain) which eventually leads to activation of an effector caspase. A recent discovery suggests that NLR-related proteins exist in bacteria, where they recognize phage proteins and undergo ligand-induced oligomerization for activation ^41^.

Hachiman may represent another example of a bacterial defence system following the paradigm of monomeric nuclease unlocking and dimerization or oligomerization ^36^, albeit with the molecular details of activation remaining to be established. In case of Lamassu, the sensitive step of activation may readily become a target for phage countermeasures. For instance, phages could evolve decoy molecules to sequester liberated LmuA, thereby neutralizing the defence. The diversity of C-LmuA sequences observed in nature may reflect an evolutionary arms race between bacterial hosts and phages, with each innovation spurring further adaptations.

Bacterial cells often harbour multiple defence systems with certain combinations appearing together more often than expected by chance indicating functional cooperation ^42^. Lamassu collaborates with prokaryotic Argonaute in the restriction of plasmids in *V. cholerae* ^10^. Based on our findings, it seems unlikely that the DNA double strand breaks generated by Lamassu are used as substrates or signals by prokaryotic Argonaute directly. While DNA repair pathways could convert DSBs into ssDNA or short oligonucleotides for prokaryotic Argonaute, these however would originate from the MGE as well as host DNA, thus rendering the sequence specificity of Argonaute pointless. More likely, prokaryotic Argonaute generates molecular signatures that activate Lamassu. Conceivably, Lamassu serves to escalate the response of other defence systems, such as the one by prokaryotic Argonaute, converting a response targeted against the MGE to a general response. Consistent with this idea, UV-induced DNA damage appears to also activate the LmuACB^DdmABC^ Lamassu system in *V. cholerae*, rendering pandemic strains that carry this system more susceptible to UV-mediated cell death ^24^.

LmuB and Rad50 proteins share an overall architecture, suggesting they evolved from a common ancestor. We noticed that some LmuB proteins more closely resemble Rad50 proteins by size. Specifically, MC1-type LmuC proteins are associated with larger LmuB proteins (∼800-1000 aa), while LmuB proteins otherwise tend to be smaller (∼600 aa) than Rad50 due to shorter coiled coils ^11^. Based on structure prediction, LmuC^MC^^1^ proteins moreover share similarity with a carboxyterminal region of the Rad50-associated nuclease Mre11, including the capping domain and the Rad50 coiled coil binding domain, but not the nuclease domain itself. Accordingly, the Lamassu defence systems might be classified into two sub-families: one more closely related to Rad50/Mre11 DNA repair complexes (with LmuA^CTD^^1-CTD5^, Mre11-like LmuC^MC^^1^, and larger Rad50-like LmuB) and one more highly diverged sub-family (with LmuA^CTD6-CTD12^, LmuC^MC2-MC8^, and smaller LmuB). Our findings suggest that at least the second sub-family of Lamassu and Rad50/Mre11 do not follow the same principle, despite their structural similarities including the transitioning between collapsed and open forms. In Rad50, DNA processing is strictly limited to the immediate vicinity of the recognized DNA end, due to the stable binding of the Mre11 nuclease to Rad50 (Fig. 7B). In Lamassu, the LmuA effector appears to be liberated from LmuB-LmuC, and the recognized substrate; it therefore seems likely that active LmuA will be free to cleave any DNA. Moreover, ATP-head engagement might be sufficient for Lamassu nuclease unlocking, while Mre11 nuclease activation in Rad50 also requires the step of ATP hydrolysis ^18^. To address whether the low ATP hydrolysis activity of LmuB is critical or even relevant for phage defence it would be necessary to generate mutations (other than the usual E-to-Q mutation) that specifically block the step of ATP hydrolysis in LmuB.

Our findings not only elucidate the molecular mechanism of Lamassu activation but also raise broader questions about bacterial defence mechanisms. Could activation by unlocking and oligomerization be a common theme across other systems? How might phages counteract such mechanisms, and what strategies have bacteria evolved in response? Future research will undoubtedly build on this work, deepening our understanding of the molecular choreography between bacteria and MGEs.

## Materials and Methods

### General methods for *in vivo* assays

The *E. coli* strains, *V. cholerae* strains, bacteriophages used for *in vivo* assays, and plasmids for cellular experiments are summarised in Table S2, S3, S4, and S5, respectively. All *V. cholerae* strains are derivatives of the 7th pandemic O1 El Tor strain E7946 ^43^. *E. coli* strain MG1655 was used for *Escherichia* phage plaque assays. *E. coli* strains S17-1 A-pir ^44^ and MFDpir ^45^ were used for plasmid maintenance and for bacterial mating with plasmids carrying the conditional R6K origin of replication. Bacterial strains were cultured at 37°C in Lysogeny broth (LB-Miller; 10 g/L tryptone, 5 g/L yeast extract, 10 g/L NaCl; Carl Roth, Switzerland). Where appropriate, antibiotic selection was done with Ampicillin (100μg/mL) and Gentamicin at either 50 μg/mL (*V. cholerae*) or 25 μg/mL *(E. coli*). Genes under the control of the arabinose-inducible *P*BAD promoter were induced by the inclusion of L-arabinose in the growth media at either 0.02 or 0.2%, as indicated.

### Strain construction

Scar-less and marker-less deletion of *V. cholerae lmuACB^ddmABC^*was performed by allelic exchange using the counter-selectable plasmid p28τι*lmuACB^ddmABC^*^46^ delivered via biparental mating from *E. coli*, as previously described ^47^. Genes encoding *lmuACB^ddmABC^* homologues from diverse organisms (Table S1) were codon-optimised for expression in *E. coli* and synthesised by Twist Bioscience, USA. Systems were then amplified by PCR and cloned into plasmid pGP704-TnAraC, which harbours a mini-Tn7 carrying *araC* and the *araBAD* promoter (*P*BAD) ^48^. pGP704-TnAraC derivatives carrying the gene(s) of interest under the control of the arabinose-inducible *P*BAD promoter were integrated into a neutral locus downstream of *glmS* in *E. coli* and *V. cholerae* strains by tri-parental mating, as previously described ^49^. Plasmids were introduced into chemically-competent *E. coli* cells using a standard heat-shock protocol. Site-directed mutagenesis of plasmid DNA was done by inverse PCR using Q5 High Fidelity DNA Polymerase (NEB, USA). All constructions were verified by PCR and either Sanger sequencing or Oxford Nanopore Technologies (ONT)-based full plasmid sequencing (Microsynth AG, Switzerland).

### *E. coli* bacteriophage plaque assays

Overnight cultures of *E. coli* MG1655 strains containing chromosomally integrated, arabinose-inducible versions of the various LmuACB^DdmABC^ homologues were back-diluted 1:100 in 3 mL LB supplemented with 5 mM MgCl2, 5 mM CaCl2, and 0.2% arabinose, and grown at 37°C, 180 rpm for 2h to exponential phase. Cultures were then diluted 1:40 in molten top agar (LB + 0.5% agar) supplemented with 5 mM MgCl2, 5 mM CaCl2, and 0.2% arabinose, poured onto a LB + 1.5% bottom agar base, and allowed to solidify at RT for 1h. Due to toxicity, cultures producing *SalEnt* LmuACB were diluted 1:10 in top agar. Phages were serially diluted and 5 μL of each dilution spotted out onto the seeded plates. Plates were imaged and enumerated after incubation overnight at 37°C. Fold-protection is expressed as the ratio between the number of plaque forming units (PFU) present on the no system control strain (MG1655-TnAraC) compared to the number of PFU present on the strains producing the various Lamassu homologues, and is expressed as the mean of three independent biological repeats.

### *V. cholerae* bacteriophage plaque assays

*V. cholerae* strains were grown overnight at 37°C, 180 rpm in 3 mL LB supplemented with 0.02% arabinose. 50-100 μL of overnight culture was then added to 3.5 mL molten top agar, poured onto a LB + 1.5% bottom agar base, and allowed to solidify at RT for 30 min, before 4 μL of a serial dilution of vibriophage X29 was spotted onto the seeded plates. Plates were imaged after incubation overnight at 37°C, the PFU/mL enumerated, and data are shown as the mean of three independent biological repeats. The detection limit in this assay was 2.5×10^3^ PFU/mL.

### Chimeric cJun fusions and toxicity assays in *E. coli*

The N-terminal domain aa 1-254 of *S. enterica* LmuA (*n*-*lmuA*) was fused to the cJun leucine zipper motif (RIARLEEKVKTLKAQNSELASTANMLREQVAQLKQKVMNH) either without (*n*-*lmuA*-*cJun*) or with (n-*lmuA*-*GSG*-*cJun*) a GSG linker. The N-terminal domain aa 1-239 of *V. cholerae* LmuA^DdmA^ was fused to the cJun leucine zipper motif with either a single alanine linker (*n*-*lmuA^ddmA^*-*cJun*) or with a GSG linker (*n*-*lmuA^ddmA^*-*GSG*-*cJun*). Overnight cultures of *E. coli* S17-1 1*pir* carrying pGP704-TnAraC encoding the various constructs were serially diluted, 5 μL of each dilution spotted onto LB plates in the absence and presence of 0.02% or 0.2% arabinose, and toxicity evaluated after incubation overnight at 37°C. Images shown are representative of three independent biological repeats.

### Toxicity assays in *V. cholerae*

Overnight cultures of *V. cholerae* strains containing chromosomally integrated *P*BAD-*lmuACB* derivatives were serially diluted, 5 μL of each dilution spotted onto LB plates in the absence and presence of 0.2% arabinose, and toxicity evaluated after incubation overnight at 37°C. Images shown are representative of three independent biological repeats.

### Protein expression and purification

*SalEnt* LmuACB and mutant variants were cloned into vectors as described in Table S5 by 4G cloning ^29^. Note that pLIBT7 vectors were used analogous to pMulti vectors for single gene expression. N-terminal 6xHis-GST-tev tagged LmuA was co-expressed with LmuB and LmuC in *E. coli* BL21(*DE3*). Cells were grown at 37°C until OD600nm=0.6 in LB medium supplemented with antibiotics and then shifted to 16 °C for 16 h; IPTG (150 µM final) was added to induce protein expression. Cells were harvested and washed once with PBS buffer, and resuspended in GST buffer (300 mM NaCl, 50 mM Tris pH 7.5, 5 mM 1-mercaptoethanol 5% glycerol) containing protease inhibitor cocktail. Cells were lysed by sonication on ice with a VS70T tip using a SonoPuls unit (Bandelin), at 40% output for 20 min with pulsing (5 s on / 5 s of). The lysates were centrifuged at 16000 rpm for 1h using a JLA-16.250 rotor (Beckman). The supernatant was loaded onto a 5 ml Glutathione Sepharose 4 Fast Flow resin (GE Healthcare) using a peristaltic pump (ISMATEC) and washed with 50 mL GST buffer and then eluted by adding 10 mM reduced L-Glutathione (Sigma-Aldrich) into GST buffer. The HisGST tag was cleaved by TEV during overnight dialysis at 4 °C in dialysis buffer (200 mM NaCl, 50 mM Tris pH 7.5, 5 mM 1-mercaptoethanol). Cleaved protein was diluted into 100 mM NaCl the next day and applied onto a 5 ml Heparin column (Cytiva) and eluted via a linear gradient of buffer B (1 M NaCl, 50 mM Tris pH 7.5, 5 mM 1-mercaptoethanol), and further purified using a Superdex 200 Increase 10/300 GL column (Cytiva) in GF buffer (150 mM NaCl, 20 mM Hepes-KOH pH 7.5 1 mM tris(2-carboxyethyl)phosphine, TCEP). The final purified proteins were concentrated using a 10 kDa cutoff concentrator (Sartorius) and flash-frozen in liquid N2 for storage at −70°C.

To isolate LmuA, the same HisGST-TEV tagged LmuA was co-expressed with LmuC and an C-terminal uncleavable His tagged LmuB (G318R). Purification was performed as with LmuACB, but with an additional high salt (1.5 M NaCl) GST buffer wash during the GST bead washing step. Fractions collected after Heparin elution were further purified with Superdex 200 Increase 10/300 GL column (Cytiva) in GF buffer.

### Cryo-electron microscopy (cryo-EM)—Data collection Structure of LmuACB^apo^

6 µM of fresh LmuA^KA^CB were reconstituted in buffer N150 (150mM NaCl, 100mM Hepes pH 7.5, 1mM TCEP, 5mM MgCl2). The complexes were supplemented with 0.05% (v/v) n-octyl-beta-D-glucopyranoside (1-OG, ChemCruz) and 1mM ATP and incubated at room temperature for 10 min. Quantifoils R1.2/1.3 mesh 400 grids were freshly glow discharged in an EasyGlow device with 15 mA current for 30 s. 3 µl of LmuABC was then applied to the grids and flash frozen in liquid ethane using a Thermo Fisher Scientific Vitrobot IV (4 s blotting time--10 blotting force) using Whatman filter paper 1. Cryo-EM images were collected on Thermo Fisher Scientific Titan Krios microscope operating at 300 keV using a Falcon 4i (TFS) camera with 96000 x magnification, yielding pixel sizes of 0.83 Å/pixel and a total dose of 50 electrons. 15817 movies were collected in total and saved in the EER format using the EPU software, defocus range from −0.8 to −2 µm (Extended Data Fig. 8A).

### Structure of LmuACB^DNA^

10 µM of LmuA^KA^CB and a 40-mer DNA duplex (Table S7) were mixed and incubated at room temperature for 30 min in buffer N50 (50 mM NaCl, 20 mM Hepes pH 7.5, 1 mM TCEP, 5 mM MgCl2) samples were further gel-filtrated with Superose 6 increase 10/300 size exclusion chromatography (SEC) column (Cytiva) Compared to the LmuA^KA^CB^apo^, the shifted peak that is supposed to be DNA bound complex were collected and concentrated to A280 is around9. 1-OG (0.05% v/v) and 1 mM ATP were then supplemented and incubated on ice before grid freezing. Cryo-EM images were collected on Thermo Fisher Scientific Titan Krios microscope operating at 300 keV using a Falcon 4i (TFS) camera with 165000 x magnification, yielding pixel sizes of 0.83 Å/pixel and a total dose of 50 electrons. 22437 movies were collected with the addition of a 30° tilted dataset due to orientation bias (Extended Data Fig. 10B).

### Structure of LmuA^tetra^

40 µM of LmuA (see above) were mixed with 10µM 45bp dsDNA (Table S7) in buffer K50 (50 mM KCl, 20 mM Hepes pH7.5, 5 mM MgCl2,1 mM TCEP), 0.05% 1-OG and 1mM ATP were added and incubated at 37 °C for 3 min. The grid was prepared using chameleon® (SPT Labtech) using a standard workflow protocol. The original chameleon® foil grid (self-wicking copper 300mesh R1.2/0.8) was loaded to chameleon® and grow discharged at 12 mA for 60 s. The sample was sprayed to the grid at humidity 82%, temperature 23 °C and plunge frozen within 304 ms. Settings of data collection were the same as with the other DNA binding structures, 11186 movies were collected on Thermo Fisher Scientific Titan Krios microscope operating at 300 keV using a Falcon 4i (TFS) camera (Extended Data Fig. 11A).

### Cryo-electron microscopy (cryo-EM)—Data processing

Initial data processing for all structures was performed on-the-fly using CryoSPARC Live ^50^, followed by particle picking and 2D classification. Dose-weighted micrographs, collected as described in Extended Data Fig. 10–11, were used as input after CTF estimation and blob picking. Subsequent steps included multiple rounds of 2D classification and ab-initio reconstruction. The initial volumes were employed for template-based particle picking, with further refinement steps outlined in Extended Data Fig. 10–11.

For the LmuACB DNA-binding structure, additional classification using 3D variability analysis ^51^ and 3D variability display (cluster mode) was performed to better resolve the distinct DNA-binding ends. The same approach was applied to the LmuA tetramer, as the CTD tetrameric domain exhibited significant mobility. Final volumes were generated using unbinned particles.

Model building was initiated using AF2 or AF3 predictions for individual subunits or the complex ^52,53^. These models were then fitted into the cryo-EM density maps using UCSF ChimeraX ^54^, followed by manual adjustments in Coot v0.9.6 ^55^ and ISOLDE ^56^. Model refinement and validation were performed using Phenix v1.19.2 ^57^.

### DNA end protection assay

2 nM 5 kb linear dsDNA with a single EcoRI cleavage site was incubated in 15 μL reactions with 2 µM LmuA^K68A^CB at RT for 15 min in in buffer N50 (containing 1 mM ATP if needed). T5 exonuclease (2 U) or EcoRI (4 U) were then added and incubated at 37°C for 10 min, the reaction was stopped by adding 3 μL 6 x SDS loading dye (Thermo Fisher Scientific R1151) and heated at 70°C for 10 min. 16.5 µL of the reaction were loaded and run in a 1% (w/v) agarose gel stained with ethidium bromide (EtBr). Electrophoresis was then performed at 5 V/cm for 1 hour and bands visualized with a transilluminator (UVP GelSolo).

### Fluorescence anisotropy assay

40-mer3’ 6-FAM labelled dsDNA was prepared by annealing two complementary DNA strands (Table S7) in buffer (50 mM Tris pH 7.5, 50 mM NaCl). Fluorescence polarisation assays were conducted in a buffer containing 50 mM Tris pH 7.5, 50 mM NaCl, 5 mM MgCl2, 0.5 mM TCEP and 1 mM ATP if needed. A series of protein concentrations, ranging from 0-2 µM, were incubated in the presence of 50 nM DNA for 30 min at room temperature to attain equilibrium. Fluorescence polarization was recorded immediately thereafter using a Synergy Neo Hybrid Multi-Mode Microplate reader (BioTek) with the appropriate filters in black 96-well flat bottom plates at 25 °C.

Data was analyzed and plotted with GraphPad Prism 10 software.

### Nuclease assay

Nuclease assays were performed with 1.8 kb pDonor, which in most assays was negatively supercoiled obtained by QIAGEN Plasmid Maxi Kit. To prepare nicked and linear forms of pDonor, 10 μg of pDonor was incubated in 100 μL reactions with ScaI (100 U in 1x NEB rCutsmart buffer) at 37 °C or Nt.BspQI (50 U in NEB 3.1 buffer) at 50 °C respectively for 1 hour, followed by column purification.

In 15 μL reactions, pDonor (8.5 nM final) was mixed with LmuACB (typically 125 nM final, otherwise as indicated) in NEB 4 buffer supplemented with ATP (1 mM final). The reactions were incubated at 37 °C for 20 minutes, followed by addition of 3 μL 6x purple loading dye (New England Biolabs). They were then loaded on a 1% (w/v) agarose gel stained with EtBr. Electrophoresis was then performed at 5 V/cm for 1 hour and bands visualized with a transilluminator (UVP GelSolo).

Time course experiments were performed with the same reaction composition but scaled up. At indicated time points 15 μL of the reaction was transferred to chilled tubes containing 3 μL 6x purple loading dye.

For the re-ligation experiment, a 564 bp PCR product was treated with LmuACB (500 nM) for 60 minutes to fully digest the DNA. Following the reaction, the DNA was further purified and 200 ng of it was mixed with T4 ligase (Thermo Fisher Scientific, 1 Weiss Unit) in 1 x T4 ligase buffer overnight at 16 °C. The samples were run in the same conditions as above but with a 2% agarose gel.

### ATPase assay

ATPase activity measurements were done with 300 nM final SmcScpAB ^58^ and LmuACB in 100 μL pyruvate kinase/lactate dehydrogenase coupled reactions at 25°C in ATPase buffer (20 mM Hepes pH 7.5, 50 mM KCl, 5 mM MgCl2 and 1 mM TCEP). The reactions contained 1 mM NADH, 3 mM phosphoenol pyruvic acid, 100 U pyruvate kinase, 20 U lactate dehydrogenase, 1 mM ATP and 1 μM 40mer dsDNA if needed. ADP accumulation was monitored for 1 h by measuring absorbance changes at 340 nm caused by NADH oxidation in a Synergy Neo Hybrid Multi-Mode Microplate reader. Results were analysed and plotted using the GraphPad Prism 10 software.

### ENDO-Pore cleavage site mapping

ENDO-Pore cleavage site mapping was performed as described previously ^30^ with modifications as follows. Manufacturer’s protocols were used unless specified otherwise. Briefly, Lamassu treated pDonor DNA was taken at various timepoints and purified by EconoSpin® all-in-one DNA mini spin columns. Treated DNA was end-repaired using the NEBNext® Ultra™ II End Repair/dA-Tailing Module (New England Biolabs) prior to purification using a Clean and Concentrate kit from Zymo Research. Pure DNA was ligated using T4 DNA ligase (New England Biolabs) with the pUC-ori cassette then purified using AMPure beads (Beckman Coulter) prior to being used to transform MegaX DH10B™ T1R Electrocomp™ Cells (Thermo Fisher). After incubation on LB agar containing 34 µg/mL chloramphenicol at 37 °C overnight, colonies were lifted in LB and the plasmid library extracted using a HiSpeed Plasmid Midi Kit (QIAGEN). Plasmids were enriched using Rolling Circle Amplification (RCA) with EquiPhi29™ DNA Polymerase (Thermo Scientific) with the Exo-Resistant Random Primer (Thermo Scientific). RCA DNA was purified with AMPure beads before debranching with T7 Endonuclease I (New England Biolabs) using 10 U per µg DNA. Samples were incubated at 37°C for 15 minutes before the addition of 0.8 U proteinase K and a further incubation for 5 minutes at 37°C. After debranching, DNA was purified using AMPure beads before the processing with the Short Read Eliminator XS kit (PacBio).

Processed DNA was then sequenced using the Native Barcoding Kit 24 V14 (Oxford Nanopore) on an R10.4.1 flowcell for 16 hours to generate approximately 150,000-200,000 reads per sample. The resulting POD5 sequencing files were demultiplexed using Dorado V8.1 (github.com/nanoporetech/dorado) using the sup_v5.0 model, before demultiplexing using the appropriate kit. Demultiplexed sequencing FASTQ files were processed using C3POa v2.2.3 (github.com/rvolden/C3POa) using the pUC-ori cassette as the DNA splint. C3POa consensus FASTA files were further processed using the Cleavage Site Investigator (github.com/sjcross/CleavageSiteInvestigator) before analysis of the resulting output files using Excel (www.microsoft.com), WebLogo 3.7.12 (https://weblogo.threeplusone.com/) and GraphPad Prism 10.3.0 (www.graphpad.com).

## Acknowledgements

The cryo-EM grid preparation, data collection, and initial image processing were performed at the Dubochet Center for Imaging (DCI) in Lausanne (a common initiative from EPFL, UNIGE, UniBE, and UNIL). We thank Florian Roisné-Hamelin for technical help and Kyle Muir for advice during data processing. We are grateful to all members of the Gruber lab for stimulating discussions, technical help, and helpful feedback on a version of the manuscript. We thank the Félix d’Hérelle Reference Center for Bacterial Viruses from the Université Laval (Québec, Canada) for providing phage X29 (HER66). Work in the Gruber lab was supported by the SNSF project funding (10.001.017), an ERC Consolidator Grant (724482-ChroCoDyle) and internal funding. Work in the Blokesch lab was supported by an ERC Consolidator grant (724630-CholeraIndex) from the European Research Council and by EPFL intramural funding. Work in the Szczelkun lab was supported by a grant from the Biotechnology and Biological Sciences Research Council sLoLa BB/X003051/1. For the purpose of open access, a CC BY public copyright licence is applied to any Author Accepted Manuscript (AAM) version arising from this submission.

---

**Table 1.** Cryo-EM data collection, refinement, and validation statistics.

Cryo-EM data collection, refinement, and validation statistics
|  | LmuACB apo<br>PDB(9HQU)<br>EMDB(EMD-52347) | LmuACB<br>40bp dsDNA<br>PDB(9HQX)<br>EMDB(EMD-52349) | LmuA tetramer<br>45bp dsDNA<br>PDB(9HR5)<br>EMDB(EMD-52355) |
| --- | --- | --- | --- |
| **Data collection and Processing** |  |  |  |
| Microscope | TFS Titan Krios | TFS Titan Krios | TFS Titan Krios |
| Voltage(keV) | 300 | 300 | 300 |
| Camera | Falcon IVi | Falcon IVi | Falcon IVi |
| Magnification | 96Kx | 165Kx | 165Kx |
| Pixel size at detector (Å/pixel) | 0.83 | 0.726 | 0.726 |
| Total electron exposure (e<sup>-</sup>/Å<sup>2</sup>) | 50 | 50 | 50 |
| Exposure time (s) | 5.7 | 3.5 | 3.5 |
| Beam size (nm) | 580 | 500 | 500 |
| Defocus range (µm) | -0.8 to -2 | -0.8 to -2 | -0.8 to -2.2 |
| Automation software | EPU | EPU | EPU |
| Energy filter slit width (eV) | 10 | 10 | 10 |
| Micrographs collected (no.) | 15817 | 22437 | 11186 |
| Micrographs used (no.) | 15136 | 21110 | 11182 |
| Total extracted particles (no.) | 9567642 | 8139417 | 3623496 |
| **For each reconstruction:** |  |  |  |
| Final particles (no.) | 242023 | 135346 | 148952 |
| Point-group | C1 | C1 | C1 |
| Resolution (global, Å) |  |  |  |
| FSC 0.5 (unmasked/masked) | 3.6/3.5 | 3.6/3.5 | 3.9/3.7 |
| FSC 0.143 (unmasked/masked) | 3.2/3.1 | 3.1/3.0 | 3.3/3.2 |
| Resolution range (local, Å) | 1.77-30 | 2.75-30 | 2.87-30 |
| Map sharpening B factor (Å<sup>2</sup>) | -89.6 | -75.93 | -101.96 |
| Map sharpening methods | Cryosparc | Cryosparc | Cryosparc |
| **Model composition (for each model)** |  |  |  |
| Protein (residues) | 1584 | 1793 | 1264 |
| Nucleotide (nt) | 0 | 58 | 44 |
| **Model Refinement (for each model)** |  |  |  |
| Refinement package | PHENIX | PHENIX | PHENIX |
| - real or reciprocal space | real space | real space | real space |
| Model-Map scores |  |  |  |
| -CC | 0.65 | 0.69 | 0.76 |
| B factors (Å<sup>2</sup>) |  |  |  |
| Protein residues (min/max/mean) | 60.51/457.53/156.44 | 22.68/297.50/100.15 | 23.22/216.22/104.73 |
| DNA (min/max/mean) | N/A | 23.57/48.31/35.80 | 15.06/167.50/71.20 |
| R.m.s. deviations from ideal values |  |  |  |
| Bond lengths (Å) | 0.011 | 0.005 | 0.005 |
| Bond angles (°) | 1.757 | 1.036 | 0.973 |
| **Validation (for each model)** |  |  |  |
| MolProbity score | 1.55 | 2.22 | 2.29 |
| CaBLAM outliers | 1.80 | 3.13 | 2.20 |
| Clashscore | 3.43 | 14.51 | 15.25 |
| Poor rotamers (%) | 1.29 | 1.90 | 2.53 |
| C-beta deviations | 0.13 | 0.00 | 0.08 |
| Ramachandran plot |  |  |  |
| Favored (%) | 95.28 | 95.10 | 95.75 |
| Outliers (%) | 0.64 | 0.28 | 0.40 |
| Q score (contour level) | 0.4450 (0.0691) | 0.3940 (0.0384) | 0.4140 (0.0329) |

## Notes

### Competing Interest Statement

The authors have declared no competing interest.

