## Extended Data Figures for "Structure and Activation Mechanism of a Lamassu Phage Defence System": Supplemental_material_letter.pdf

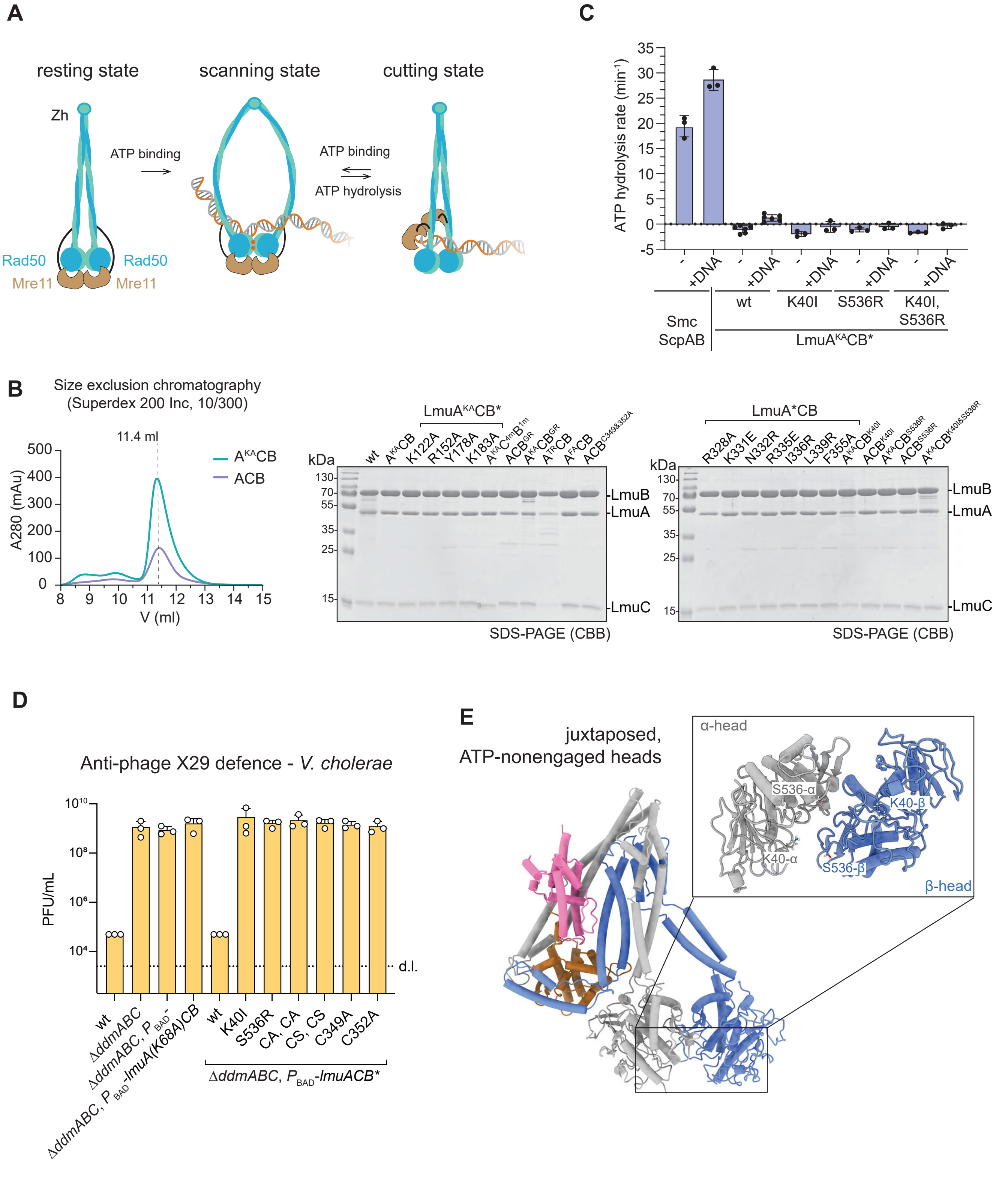


Extended Data Fig. 1

**Extended Data Fig. 1 Protein purification and characterization.**

(A) Cartoon illustration of Rad50-Mre11 scanning and cutting its DNA substrate under ATP binding & hydrolyzation.

(B) Left panel: Comparison of the gel filtration chromatography of LmuACB and LmuA$^{KA}$CB. Right panel: SDS page showing the degree of purity of preparations of LmuABC and the variants.

(C) Measurement of ATP hydrolysis activity for LmuA$^{KA}$CB and variants at 300 nM final with and without 1 μM 40-mer DNA duplex, 1 mM ATP. *Bacillus subtilis* Smc-ScpAB, as positive control, with three replicate measurements. LmuA$^{KA}$CB with six replicate measurements. LmuA$^{KA}$CB with the ATP binding mutation (K40I), ATP-dependent head dimerization mutation (S536R) and both mutations (K40I, S536R) with three replicates. Means and standard deviations are shown.

(D) Phage defence of strains with mutations of selected key residues in LmuB. As described in Fig. 1B. 'CA, CA' indicates double alanine mutation of cysteines in the Zinc hook motif; 'CS, CS', double serine mutations.

(E) Zoom view of the juxtaposed arrangement of the LmuB heads LmuACB [17]. The side chains of selection catalytic residues are indicated: ATP binding (K40) and ATP-dependent dimerization residue (S536).

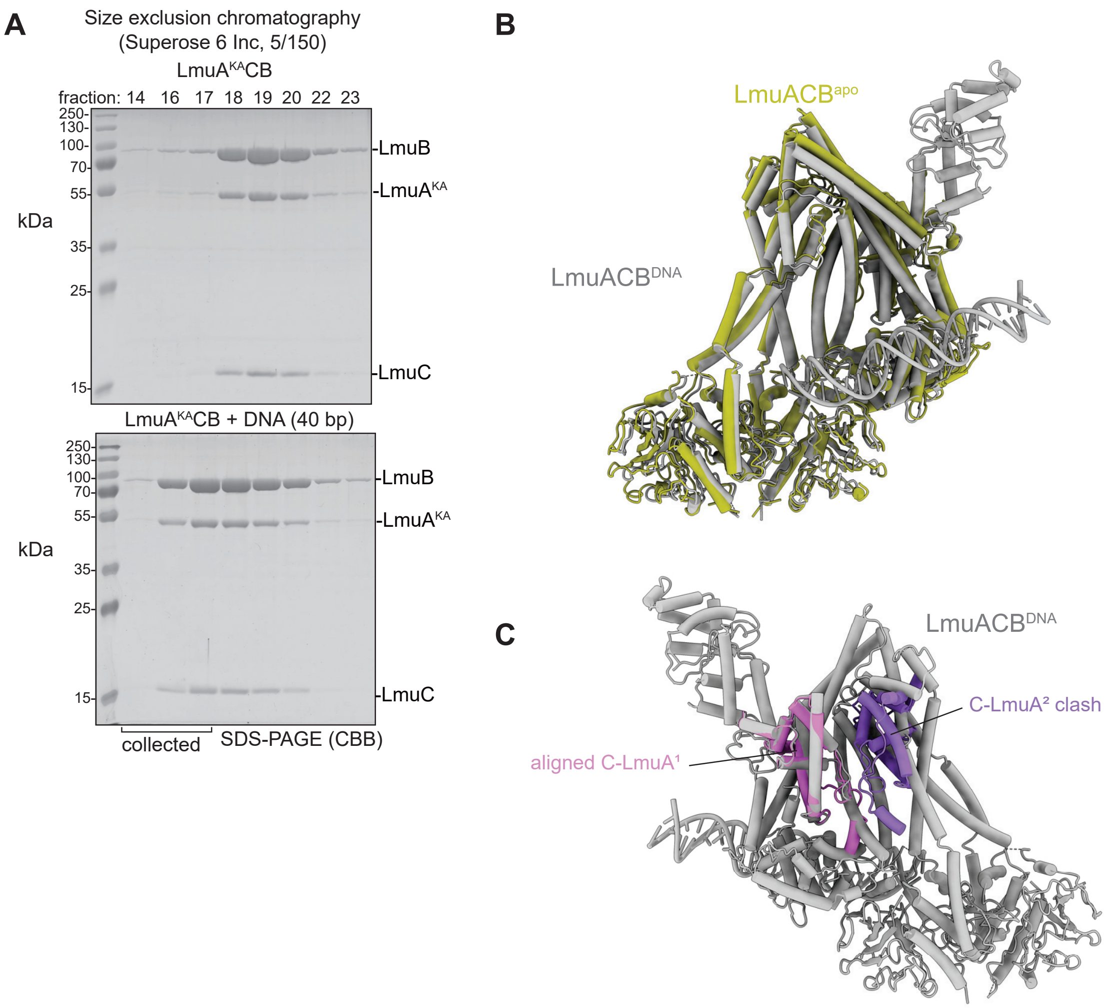


Extended Data Fig. 2

**Extended Data Fig. 2 DNA end binding.**

(A) SDS page gels showing a shift in elution from size exclusion chromatography of LmuA$^{KA}$CB with and without pre-incubation with 40-mer DNA duplex. Top gel: gel filtration fractions of LmuA$^{KA}$CB without DNA from a Superose 6 Increase 5/150 column. Bottom gel: gel filtration fractions of LmuA$^{KA}$CB preincubated with DNA.

(B) Comparinson of the LmACB$^{apo}$ (yellow) and LmuACB$^{DNA}$ (grey) model.

(C) Comparinson of sequested LmuACB$^{DNA}$ model (grey) and C-LmuA dimer (AF3). C-LmuA$^{1}$ aligned with C-LmuA from LmuACB$^{DNA}$, C-LmuA$^{2}$ shows clashes with LmuB.

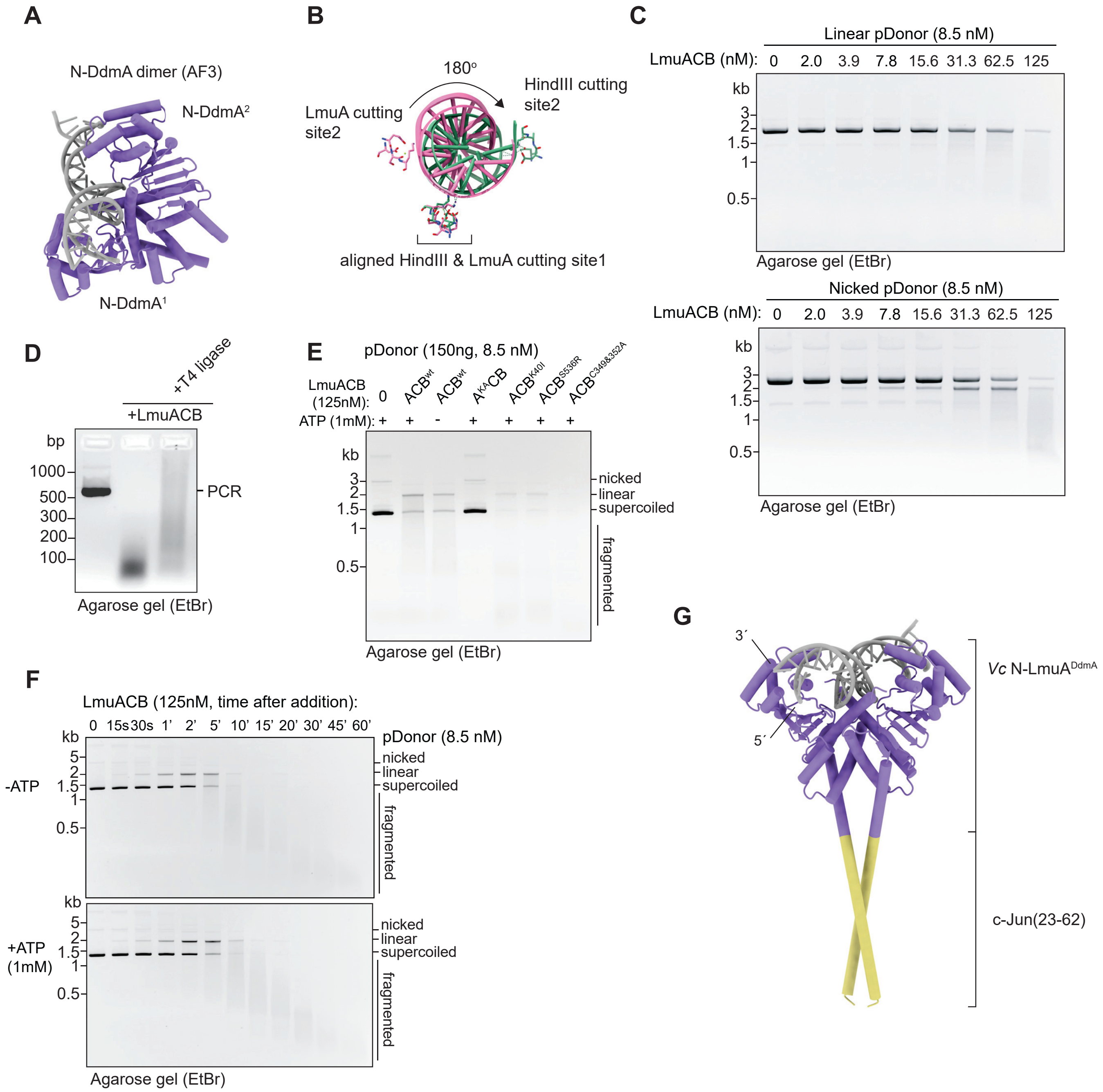


Extended Data Fig. 3

**Extended Data Fig. 3 LmuACB DNA cleavage activity.**

(A) AF3 predictions of the *V. cholerae* N-LmuA$^{DdmA}$ dimer bound to duplex DNA. As shown for N-LmuA in Fig. 3A.

(B) Direct comparison of the nuclease cutting sites in HindIII (PDB: 3WVG) and in the N-LmuA dimer (AF3). Two cutting sites (cutting site 1 in each dimer) are aligned; the relative position of cutting site 2 indicates a nearly 180° rotation between LmuA and HindIII (thus explaining the 5 nt difference in the overhangs). Catalytic residues are represented in sticks in pink (LmuA) and green (HindIII) colours.

(C) Assay for DNA degradation by LmuACB as detailed in Fig. 3B but for linearized and nicked DNA (rather than supercoiled DNA).

(D) Re-ligation of LmuACB-generated DNA fragments by T4 DNA ligase as visualized by agarose gel electrophoresis.

(E) Assay for DNA degradation by mutant LmuACB complexes. Key catalytic and structural residues in LmuB are not required for DNA degradation *in vitro*.

(F) ATP addition does not affect DNA degradation. As described in Fig. 3B but with or without ATP addition.

(G) AF3 prediction of a DNA-bound *V. cholerae* N-LmuA$^{DdmA}$ dimer fused with cJun (23-62, PDB: 2H7H). As shown for the N-LmuA dimer in Fig. 3D.

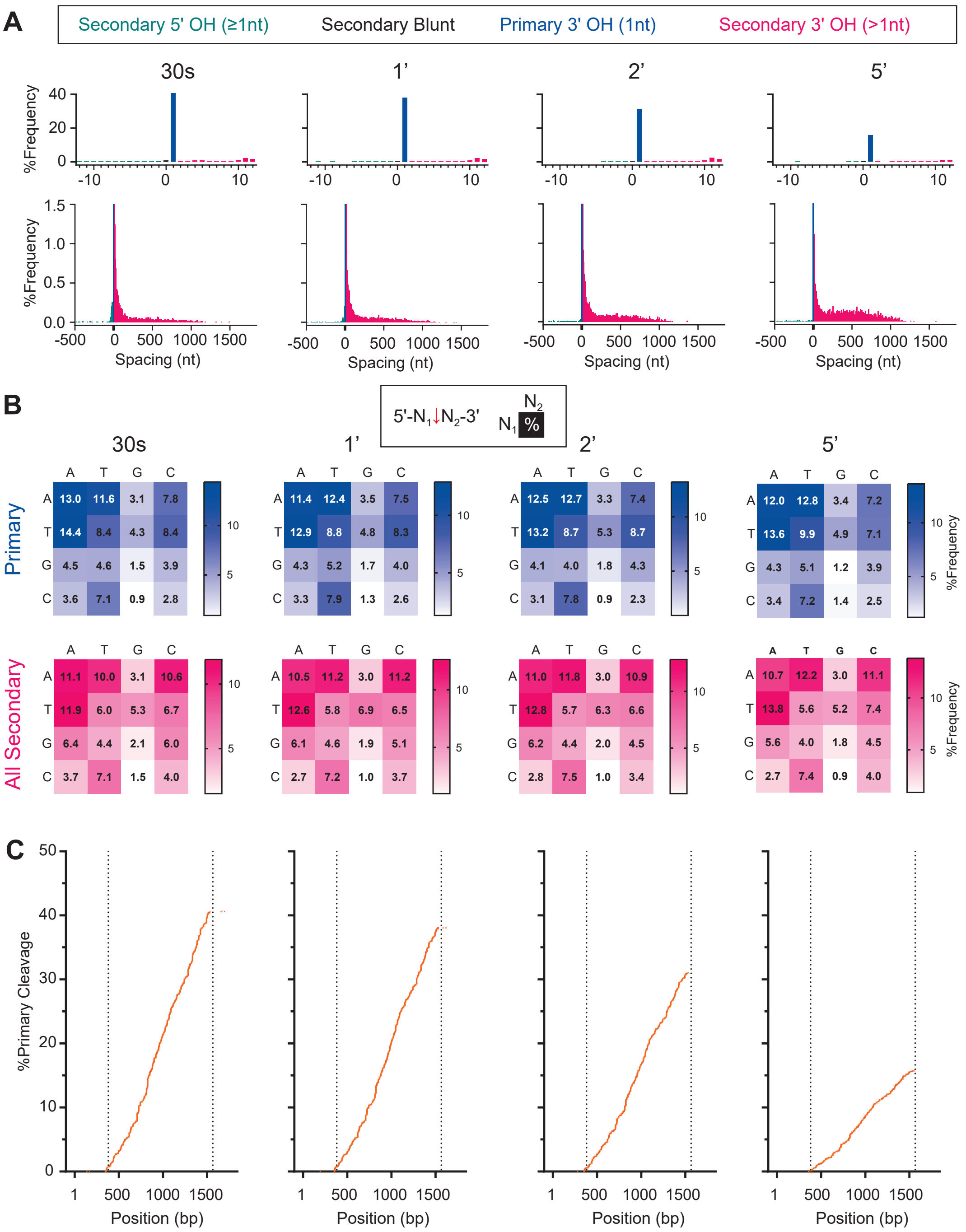


Extended Data Fig. 4

**Extended Data Fig. 4 ENDO-Pore cleavage site mapping (1).**

(A) Frequencies of primary and secondary DNA cleavage end types (blunt, 3’ overhang or 5’ overhang) at each time point following LmuACB addition as determined by ENDO-Pore sequencing.

(B) Frequencies of nucleotide preferences 5′ and 3′ of the phosphodiester cleavage site for primary and secondary events at each time point.

(C) Positional distribution of primary cleavage events shown as a cumulative frequency plot. The regions outside of the dotted lines are the chloramphenicol gene where cleavage cannot be mapped. See Extended Data Fig. 5B for the pDonor map.

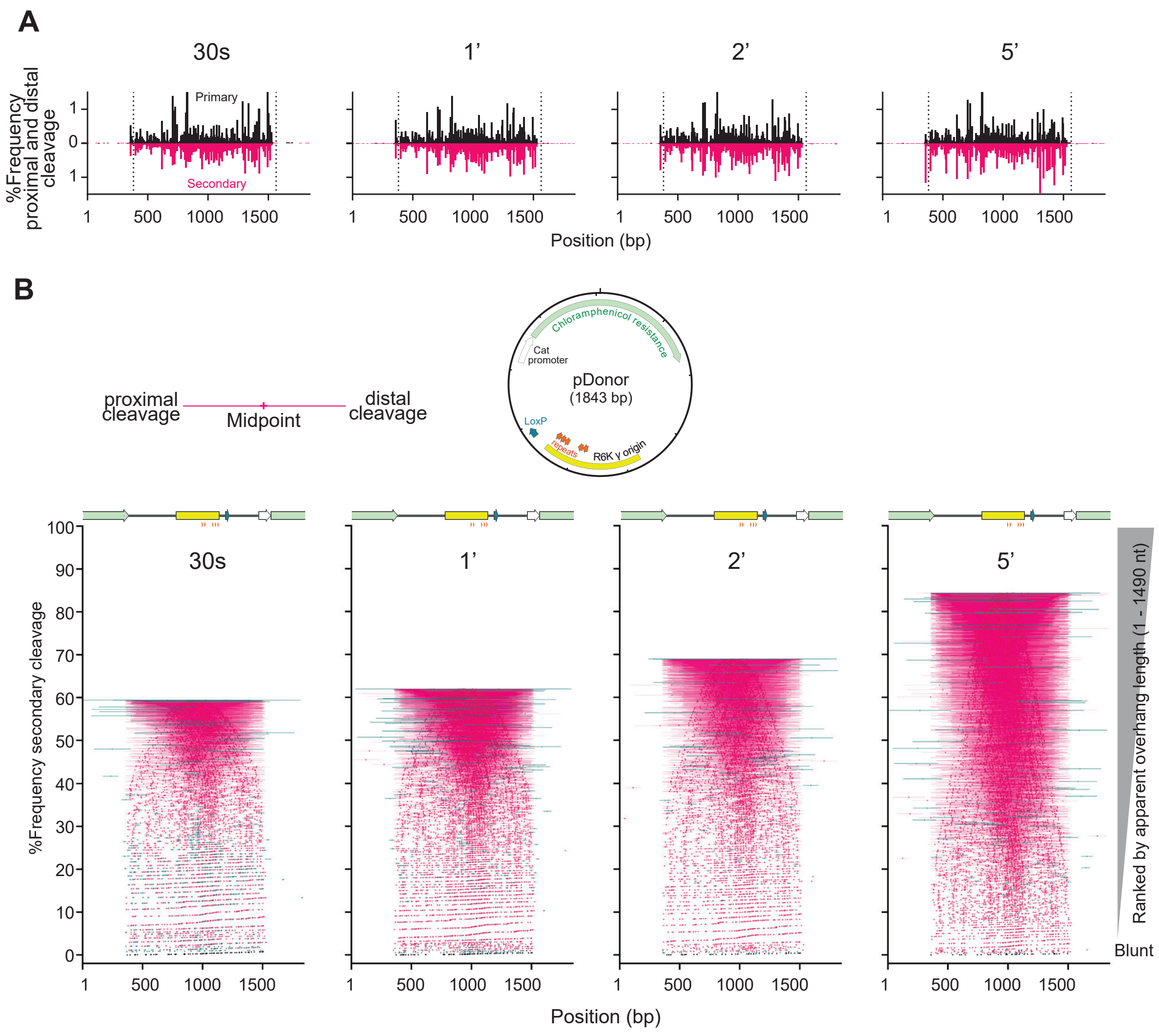


Extended Data Fig. 5

**Extended Data Fig. 5 ENDO-Pore cleavage site mapping (2).**

(A) Comparison of DNA cleavage frequencies on the top and bottom strands for primary (1 nt 3′ overhang) and secondary cleavages (all other ends) at each time point following addition of LmuACB as determined by ENDO-Pore sequencing. Hot spots in the primary cleavage positions are also observed in the secondary cleavage positions, consistent with the latter arising due to multiple cleavages starting with a primary event.

(B) Positional distribution of secondary DNA cleavages at each time point, ranked from bottom to top by apparent overhang length. Note that multiple double strand breaks in the DNA result in apparent 3′ overhangs in ENDO-Pore [30]. The long 5′ overhangs most likely result from the end-repair processing of a single strand nick and a double strand break during sequencing preparation. The cleavage midpoints are indicated by a cross. The horizontal lines indicate the spacing between cleavage sites.

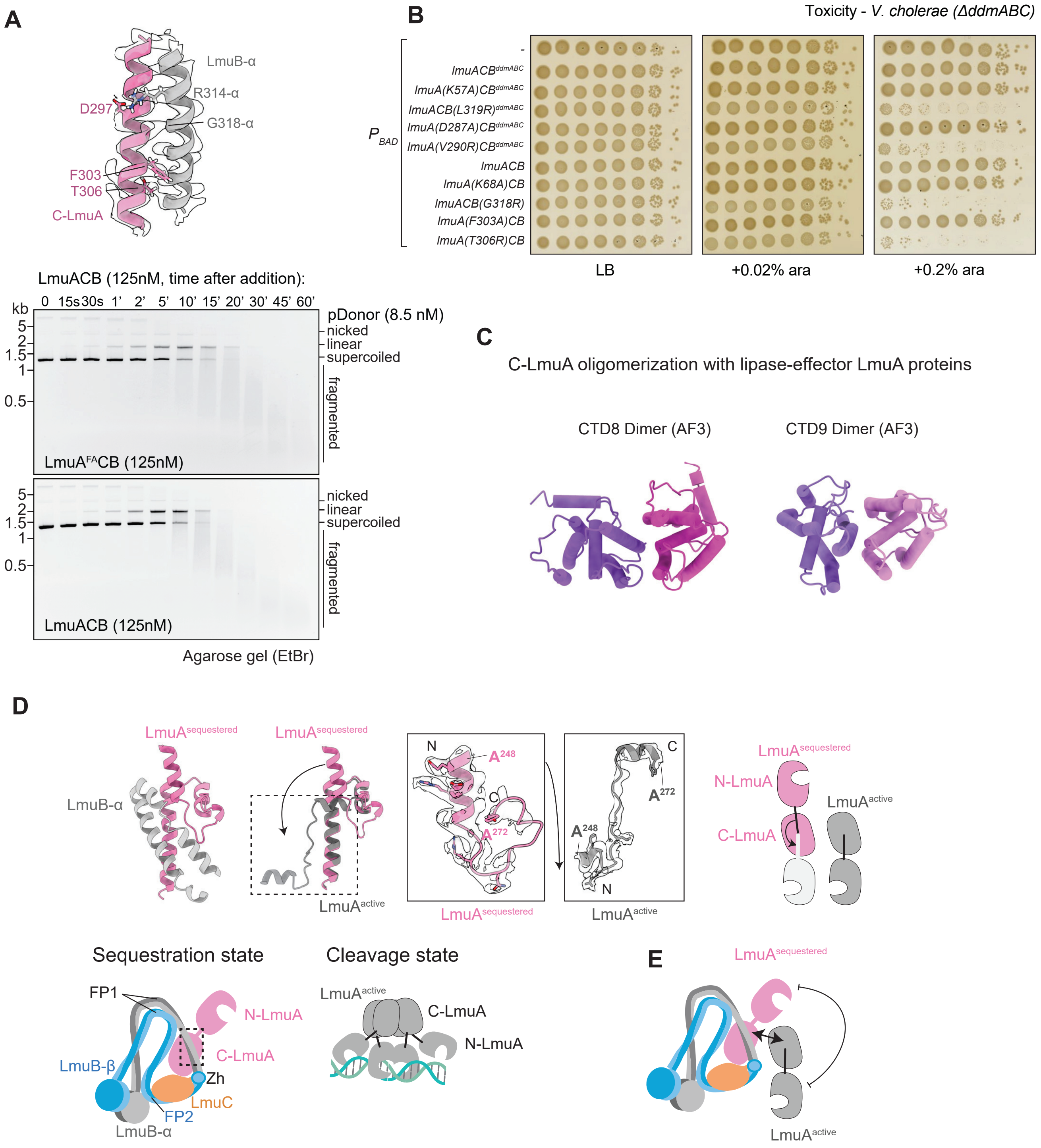


Extended Data Fig. 6

**Extended Data Fig. 6 LmuACB hyperactivity and regulation.**

(A) Top panel: Local cryo-EM map of the C-LmuA/LmuB interface, residues selected for mutagenesis (F303 and T306; notably, G318 is not visible) are indicated. Bottom panel: Agarose gel showing pDonor DNA cleavage at different time points after mixing with wild-type LmuACB (bottom image) or $LmuA^{F303A}CB$ (top image).

(B) Toxicity phenotype observed when expressing *lmuACB* mutants in *V. cholerae*, as shown in Fig. 5D but with additional strains as well as control conditions lacking arabinose (left) and having low arabinose (middle).

(C) AF3 predictions of effector oligomers of selected LmuA proteins belonging to the CTD8 (uniport: a0a061jwa6) and CTD9 (ncbi: WP_012639477) family both having a predicted lipase alpha/beta hydrolase effector domain (only dimer presented here).

(D) Comparative analysis of the N-to-C connection in LmuA. Top left: Structure of C-LmuA switching from sequestered state to active state. $LmuA^{sequestered}$ pink, LmuB-α coiled coil grey, $LmuA^{active}$ dark grey. Top middle: cryo-EM map comparison of C-LmuA $(248\text{-}272)^{sequestered}$ (pink) and C-LmuA $(248\text{-}272)^{active}$ (dark grey). Top right: cartoon illustration of N-LmuA switching nearly 180° from the sequestered state to active state relative to C-LmuA. Bottom: cartoon illustration of overall LmuA forming sequestered state together with LmuCB and switched to a tetrameric active state after being released from LmuCB, with N-LmuA turning nearly 180° compared to the sequestered state (E).

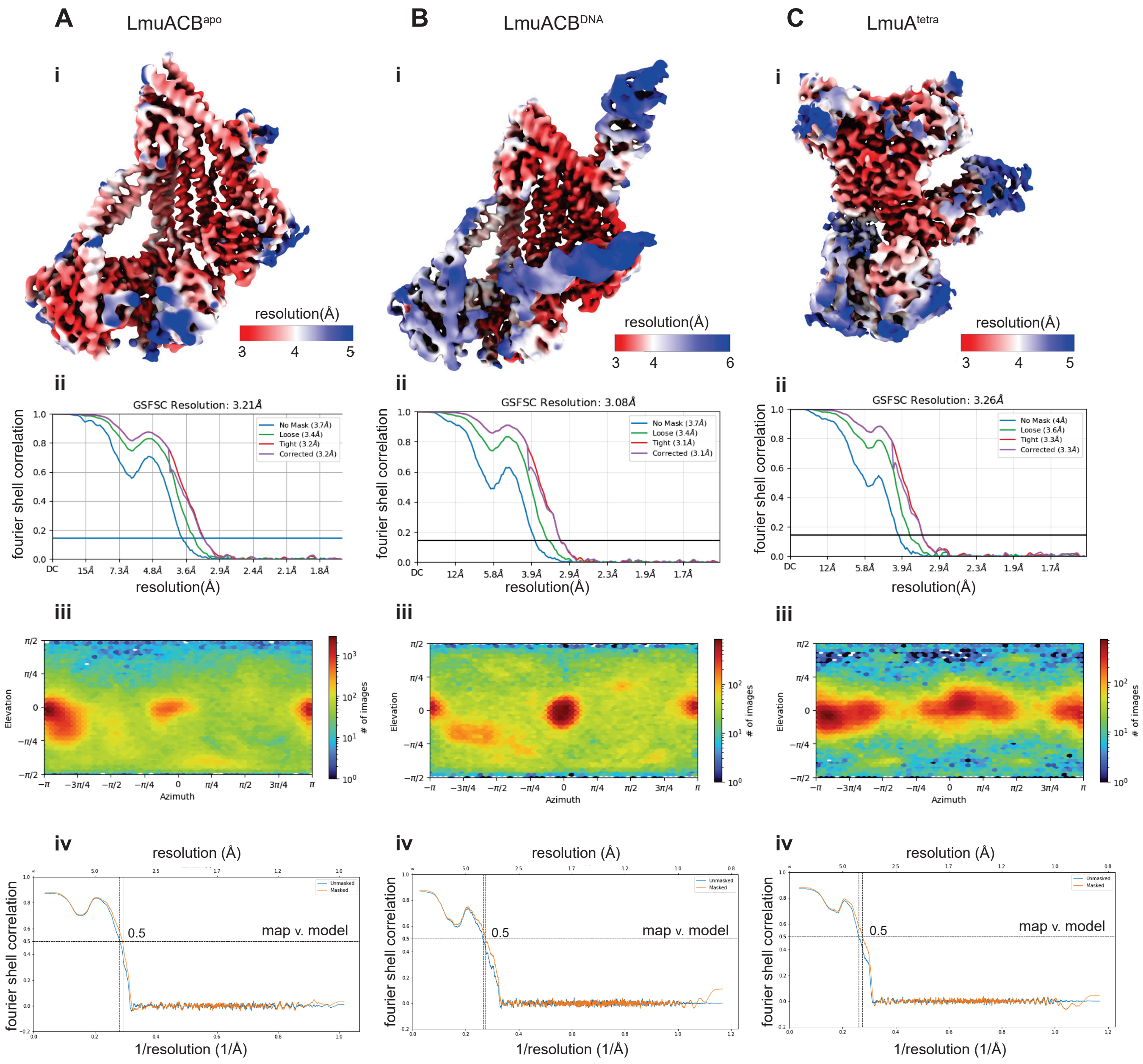


Extended Data Fig. 7

**Extended Data Fig. 7 Local resolution estimation.**

(A) – (C) Local resolution estimation for the cryo-EM structures (ChimeraX) discussed in the paper (**i**). GSFSC curves (**ii**) and Euler plots (**iii**) indicating the overall resolution and orientation distribution (CryoSparc v3&4) are shown too. Map-model FSC curves with and without mask were calculated using Phenix 1.19.2 (**iv**).

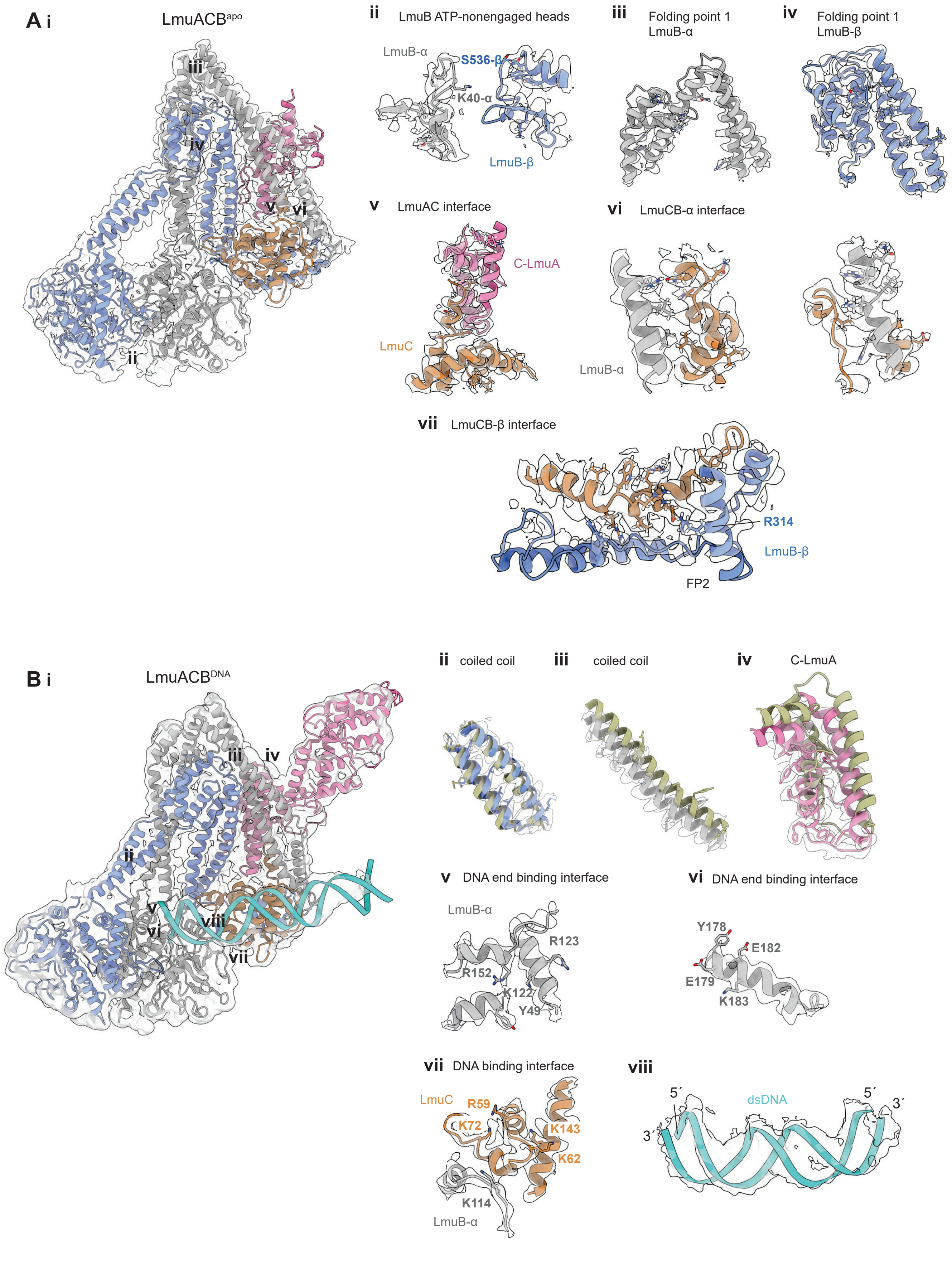


Extended Data Fig. 8

**Extended Data Fig. 8 Map-model fitting of the LmuACB$^{apo}$ and LmuACB$^{DNA}$ structures.**

(A) Map-model fitting of the LmuACB$^{apo}$ structure. **i**, overall view of the fitting. **ii**, local fitting at the LmuB ATP-nonengaged heads, K40 & S526 side chains are highlighted. **iii-iv**, fitting at the folding point 1. **v-viii**, details of LmuC extensively binding to LmuA and LmuB.

(B) Map-model fitting of the LmuACB$^{DNA}$ structure. **i**, overall view of the fitting. **ii-iv**, comparison of the minor conformational changes between LmuACB$^{apo}$ and LmuACB$^{DNA}$, models are aligned in ChimeraX with matchmaker. Check overall alignment in Extended Data Fig. 2B. **v-vii**, map details of LmuCB binding to the dsDNA. **Viii,** fitting of the dsDNA into the map.

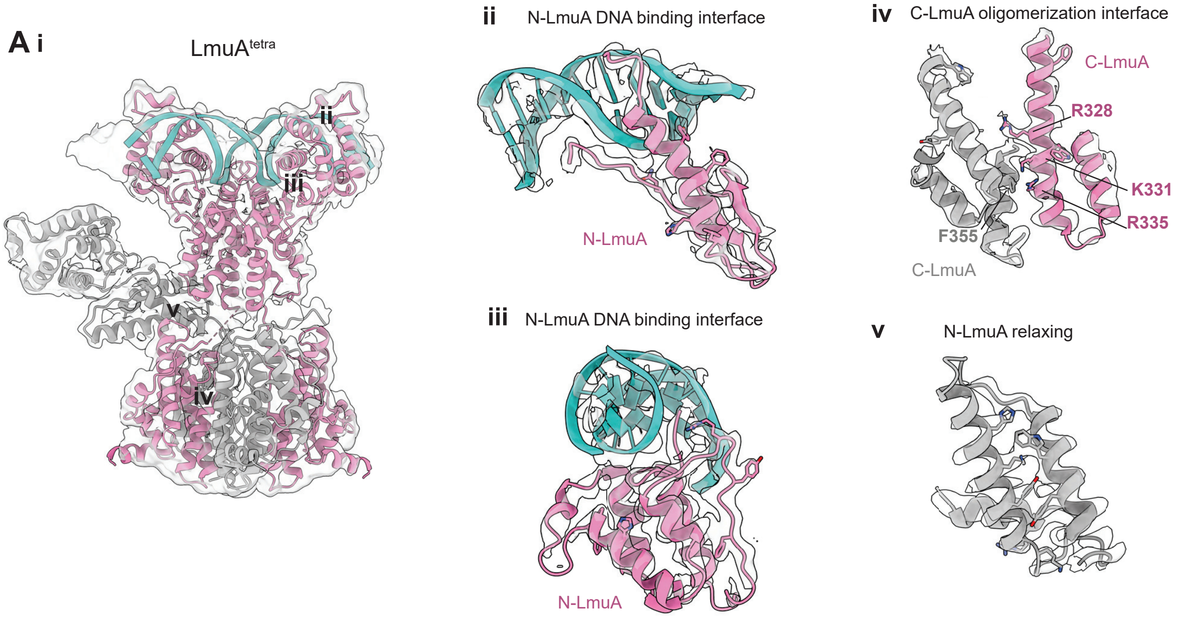


Extended Data Fig. 9

**Extended Data Fig. 9 Map-model fitting of the LmuA$^{tetra}$ structure.**

(A) **i**, overall view of the fitting. **ii-iii**, details of the N-LmuA binding to the dsDNA, some side chains are shown to prove quality of the map. **iv**, interacting interface at the C-LmuA tetramerization part. some side chains of the residues that are used previously are shown here. **v**, fitting of the relaxing N-LmuA part.

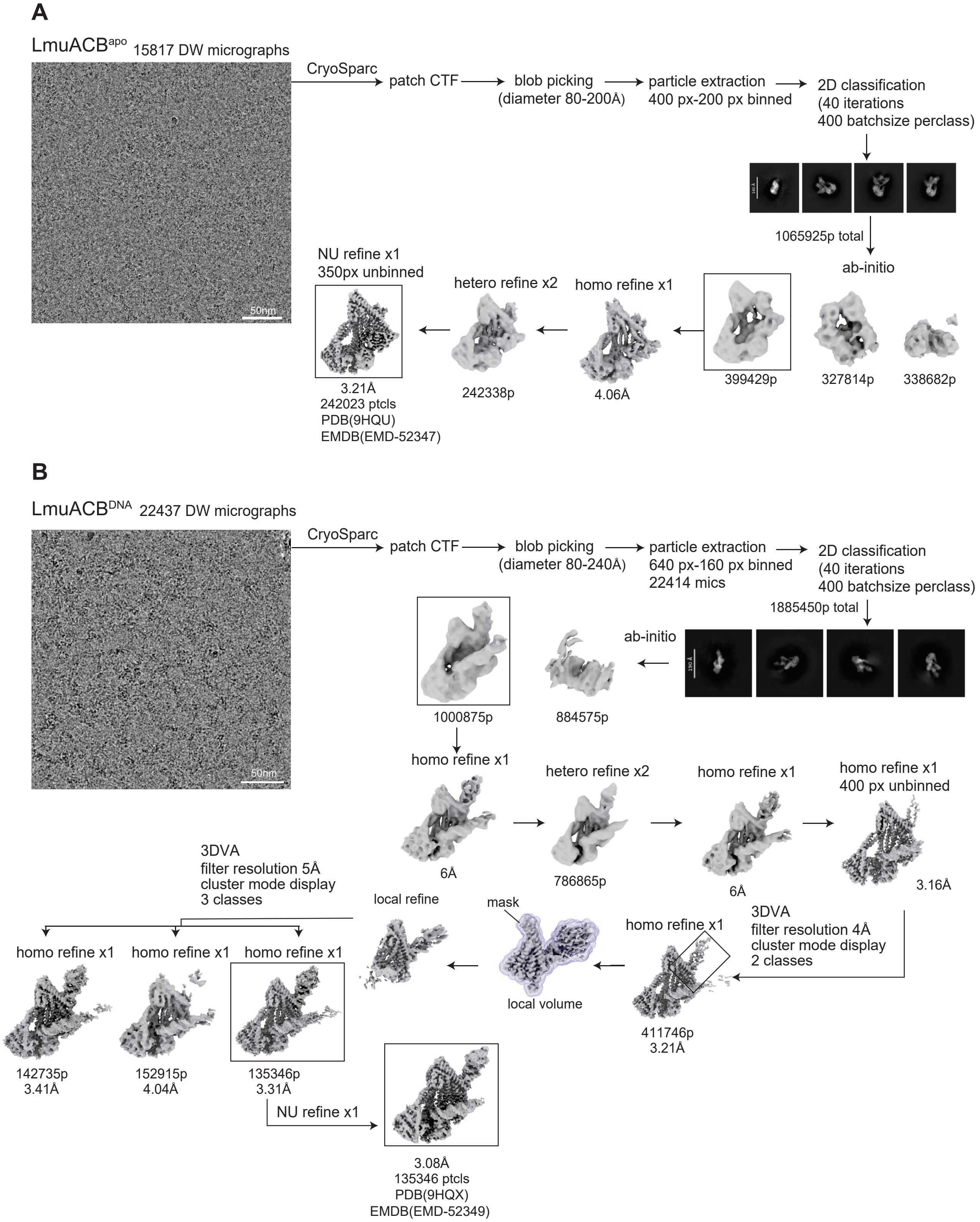


Extended Data Fig. 10

**Extended Data Fig. 10 Cryo-EM processing pipeline for the LmuACB$^{apo}$ and LmuACB$^{DNA}$ structure.**

(A) Left panel: Raw micrograph of the LmuACB$^{apo}$ structure. Right panel: processing pipeline with Cryosparc version 3.

(B) Left panel: Raw micrograph of the LmuACB$^{DNA}$ structure. Right panel: processing pipeline with Cryosparc version 3.

A

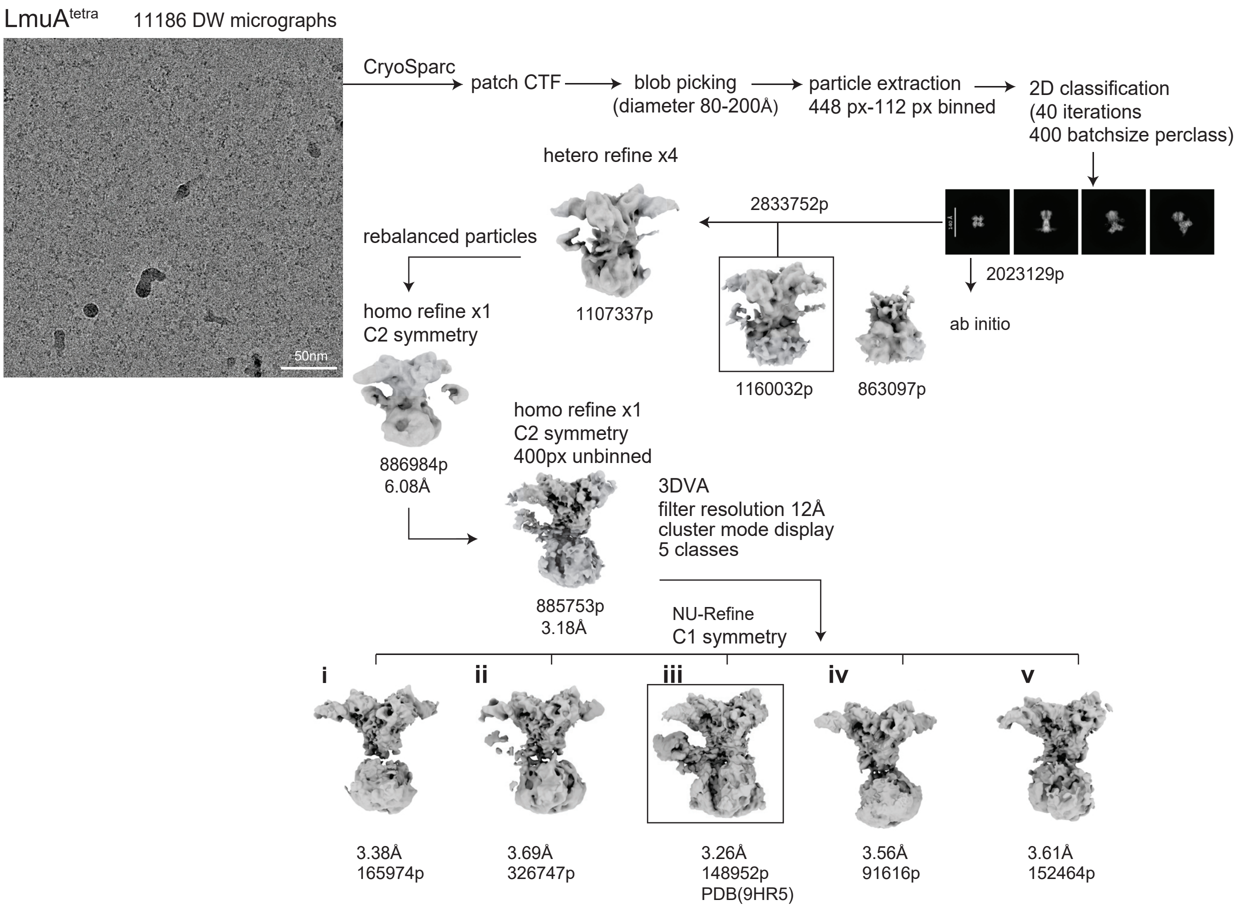


Extended Data Fig. 11

**Extended Data Fig. 11 Cryo-EM processing pipeline for the LmuA tetramer structure.**

(A) Raw micrograph of the LmuA$^{tetra}$ structure. Right panel: processing pipeline with Cryosparc version 4. Map **iii** with N-LmuA$^{relaxing}$ visible were used for determining the structure.
